# Recurrent Connectivity Shapes Spatial Coding in Hippocampal CA3 Subregions

**DOI:** 10.1101/2024.11.07.622379

**Authors:** Eunji Kong, Erfan Zabeh, Zhenrui Liao, Tiberiu S. Mihaila, Caroline Wilson, Charan Santhirasegaran, Darcy S. Peterka, Tristan Geiller, Attila Losonczy

**Affiliations:** Department of Neuroscience, Columbia University, New York, NY, United States; Mortimer B. Zuckerman Mind Brain Behavior Institute, Columbia University, New York, NY, United States; Department of Biomedical Engineering, Columbia University, New York, NY, USA; Department of Neuroscience, Yale University, New Haven, CT, USA; Wu Tsai Institute, Yale University, New Haven, CT, USA; Peter O’Donnell Jr. Brain Institute, University of Texas Southwestern Medical Center, Dallas, TX, USA; Department of Neuroscience, University of Texas Southwestern Medical Center, Dallas, TX, USA

## Abstract

Stable and flexible neural representations of space in the hippocampus are crucial for navigating complex environments. However, how these distinct representations emerge from the underlying local circuit architecture remains unknown. Using two-photon imaging of CA3 subareas during active behavior, we reveal opposing coding strategies within specific CA3 subregions, with proximal neurons demonstrating stable and generalized representations and distal neurons showing dynamic and context-specific activity. To test the causal role of excitatory connectivity in neural computation, we employed a genetic manipulation approach in which local disruption of glutamatergic synaptic transmission impaired context-specific spatial coding in distal CA3. Aligned with these experimental results, we show in artificial neural network models that varying the recurrence level causes these differences in coding properties to emerge. We confirmed the contribution of recurrent connectivity to functional heterogeneity by characterizing the representational geometry of neural recordings and comparing it with theoretical predictions of neural manifold dimensionality. Our results indicate that local circuit organization, particularly recurrent connectivity among excitatory neurons, plays a key role in shaping complementary spatial representations within the hippocampus.

**Highlights:**

- Proximodistal CA3 subregions implement complementary coding strategies in relation to time and context
- Disrupting excitatory recurrence in distal CA3 impairs context-discriminative neural coding
- Sparse and dense neural networks capture the functional heterogeneity of CA3 subcircuits
- Recurrence level tunes the geometry of neural manifolds both *in vivo* and *in silico*

## Introduction

Recurrent connectivity among excitatory neurons is a critical feature of the cortical microcircuit that underlies functions such as sensory processing, working memory, and decision-making^1-6^. In the hippocampus, strong recurrent excitatory connections among pyramidal neurons (PNs) within the CA3 region are thought to provide a basis for memory-guided behaviors^7-10^. This circuitry is believed to endow CA3 with the ability to rapidly store patterns of integrated sensory information for the initial encoding of contextual representations, and to retrieve them later from partial or degraded inputs^11,12^. However, the density of recurrent collaterals changes on a gradient along the transverse axis, from low recurrence in proximal CA3 (pCA3; close to the dentate gyrus) to higher recurrence in distal CA3 (dCA3; close to CA2)^13,14^, mirroring distinct molecular and physiological gradients recognized along this axis^15,16^. Given these differences, it is natural to ask whether CA3 subregions hold different roles in encoding and retrieving spatial information, and how these roles translate more broadly to learning and memory functions^17-25^.

To address this question, we performed a detailed functional characterization of pCA3 and dCA3 neurons in mice performing a spatial navigation task in virtual reality (VR)^26^. We characterized the dynamics of neural representations over time and across contexts, revealing prominent differences between the two subregions. To test whether these functional differences depend on excitatory recurrence, we locally disrupted excitatory synaptic transmission in dCA3 using a vesicular glutamate transporter 1 (VGLUT1) conditional knockout (cKO) transgenic line^27^, which resulted in impaired context-selective coding. Using recurrent neural network (RNN) and Hopfield network models, we confirmed that varying the level of recurrent connectivity between excitatory neurons is sufficient to explain these differences, demonstrating that distinct connectivity patterns in pCA3 and dCA3 lead to differential mechanisms of information processing.

We conclude that distinct subregions of CA3, shaped by their respective levels of recurrent connectivity, play complementary roles in spatial memory: dCA3 may be more involved in the flexible encoding of new contexts, while pCA3 may contribute to the stable retrieval of established memories, thus balancing adaptability and consistency in memory-guided behaviors^28-30^.

## Results

### Functional characterization of proximodistal CA3 during goal-oriented navigation

To first characterize the functional dynamics of different recurrent connections within CA3 in the dorsal hippocampus, we performed *in vivo* two-photon calcium imaging in a VR spatial navigation task^31,32^ (Fig. 1A-C, Supplementary Fig. 1A, B). We stereotactically injected Cre-dependent adeno-associated virus (AAV) into pCA3 and dCA3 regions of Grik4-Cre mice to selectively express GCaMP8s in CA3 PNs, and implanted a chronic imaging window above the hippocampus. Mice were head-fixed and trained to perform a goal-oriented learning (GOL) task^26,32^, where a spatially fixed operant reward was given in a 2-m virtual linear environment with a 2-s inter-trial interval between laps. After two weeks of training, mice successfully learned the reward zone (RZ), as reflected by anticipatory changes in animal’s velocity and licking (Fig. 1D). On each day, we observed a subset of place cells (0.13 ± 0.01 for pCA3, 0.31 ± 0.02 for dCA3), whose responses reliably tiled the entire virtual track (Supplementary Fig. 1C, G). We quantified individual place cells’ specificity, sensitivity, and spatial information and found higher proportions of place cells with lower levels of spatial information and place field specificity in dCA3 compared to pCA3 (Supplementary Fig. 1C-F). To assess differences in recurrent connectivity levels across CA3 subregions, we inferred effective recurrent connectivity from spontaneous population activity using multiple complementary approaches capturing linear correlations, conditional dependencies, lagged interactions, directed influences, and nonlinear relationships (Supplementary Fig. 2). Across all approaches, inferred connectivity was consistently less sparse in dCA3 than in pCA3, with this ordering preserved across analysis parameters.

**Figure 1.**
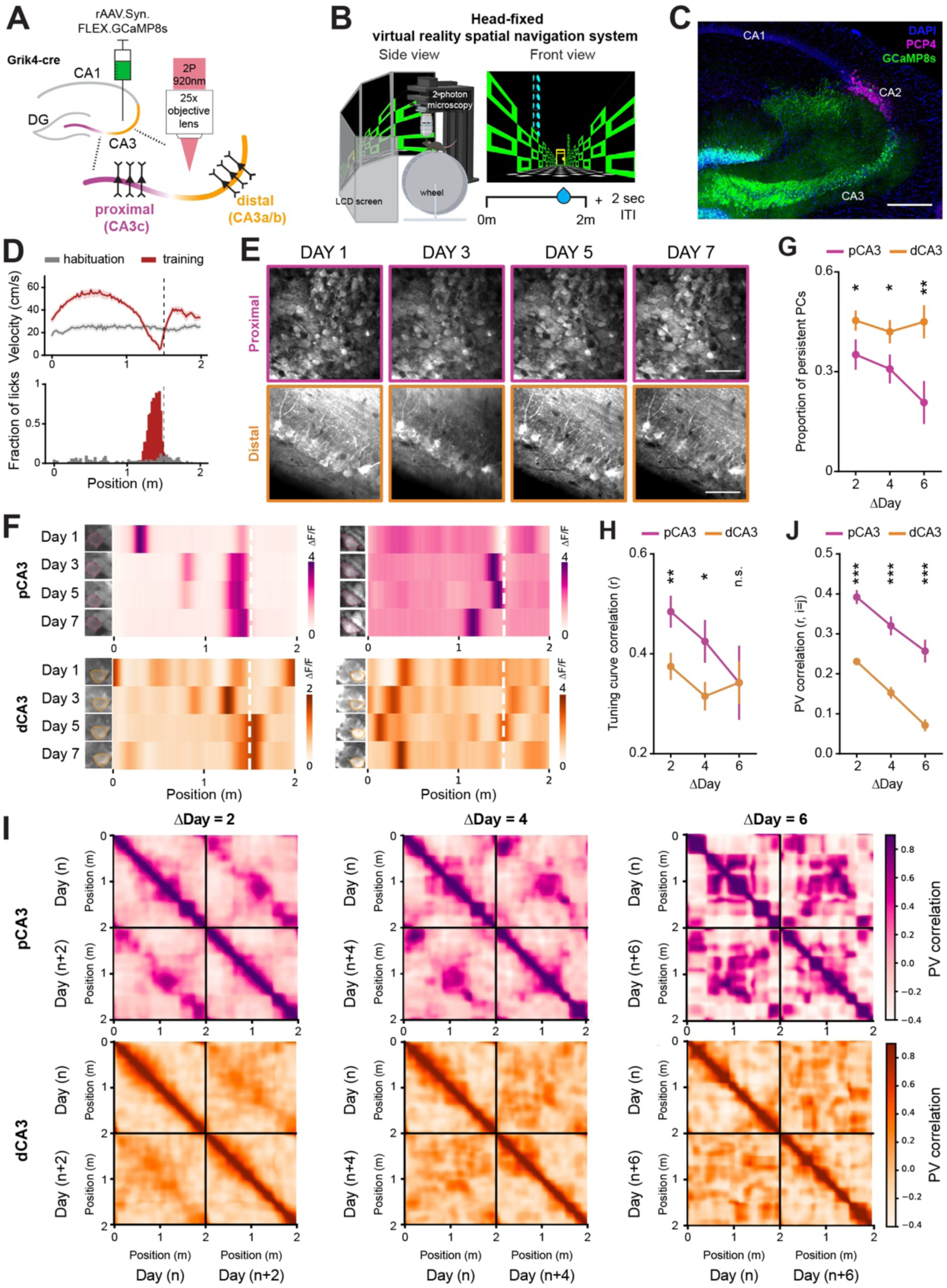
Dynamic and stable spatial representations in distal and proximal CA3. (A) Schematic of *in vivo* two-photon imaging of proximodistal CA3 neurons. Grik4-cre transgenic mice were injected with a Cre-dependent GCaMP8s virus to record the neural activities of PNs in dorsal CA3. (B) Schematic of a virtual reality navigation system combined with *in vivo* two-photon microscopy (*left*) and the task structure of a goal-oriented learning (GOL) task with a fixed reward zone in a 2m virtual environment (*right*). ITI, intertrial interval. (C) Representative confocal image showing GCaMP8s expression along the transverse axis of the CA3 region in Grik4-cre transgenic mice. Blue: DAPI, Green: GCaMP8s, Magenta: PCP4. Scale bar: 200um. (D) Examples of velocity (*top*) and licking responses (*bottom*) before and after operant conditioning of the GOL task. (E) Longitudinal and repetitive imaging of individual cells in proximal (*top*) and distal (*bottom*) CA3 subregions. Scale bar: 50 *μ*m. (F) Four examples of cross-day registered place cells and their heatmaps of spatial tuning curves with stable place fields in pCA3 (*top*) and dCA3 (*bottom*). The white dashed vertical line indicates the reward location. (G) Proportion of persistent place cells across days (mean ± SEM; two-sided unpaired t-test, p = 0.037, 0.032 and 0.0042 for ΔDay 2, 4, and 6; n = 5 and 7 mice for pCA3 and dCA3, respectively). (H) Average tuning curve correlations of place cells across days (mean ± SEM; two-sided unpaired t-test, p = 0.0092, 0.029, and 0.10 for ΔDay 2, 4, and 6; n = 5 and 7 mice for pCA3 and dCA3, respectively). (I) Heatmaps of population vector (PV) correlation between all pairs of positions across days for pCA3 (*top*) and dCA3 (*bottom*) place cells. The interval between days was 2 (*left*), 4 (*middle*), and 6 (*right*). (J) PV correlation between the same position across days for pCA3 and dCA3 place cells (mean ± SEM, two-sided unpaired t-test, p = 1.06×10^-14^, 7.17×10^-9^, and 2.80×10^-8^ for ΔDay 2, 4, and 6, n = 5 and 7 mice for pCA3 and dCA3, respectively). Colors are matched with purple for pCA3 and orange for dCA3. n.s. = non-significant, *p < 0.05, **p < 0.01, ***p < 0.001.

### Long-term dynamics of spatial representations

To investigate the long-term spatial coding properties of CA3 PNs, we tracked the same field of view every other day (Fig. 1E) as mice achieved stable behavioral performance in the GOL task. Among cross-registered cells, a subset of neurons was identified as place cells over days (Fig. 1F, Supplementary Fig. 3A, B), and we assessed whether these neurons stably encoded the same spatial location over the course of the week. We found that dCA3 place cells had a higher place cell persistence (Fig. 1G), defined as the probability that a given neuron displays a significant place field across two paired sessions, irrespective of its location on the track. This result shows that dCA3 neurons overall remain more spatially tuned across days compared to pCA3 neurons. To address whether place fields maintain their selectivity, we quantified the tuning curve correlations between sessions, and found that pCA3 place cells, though less numerous, are more stable over time than dCA3 place cells (Fig. 1H). At the population level, pCA3 population vectors remained highly correlated with each other across days compared to dCA3 population vectors (Fig. 1I, J, Supplementary Fig. 3C, D). These results imply that the dCA3 code is more dynamic and adaptable over time, while pCA3 implements a more stable, long-term code of spatial memory.

### Distal CA3 exhibits more context-discriminative neural representations

Next, to investigate the neural dynamics of proximal and distal CA3 PNs in response to contextual changes, we performed a context-switching task in which animals were exposed to a (pretrained) Familiar (F) and a Novel (N) environment in 15-lap alternating blocks (Fig. 2A, B, Supplementary Fig. 4A, B) while recording from either pCA3 or dCA3 (Fig. 2C). To quantify how contextual information is differentially processed at the single-cell level, we calculated a discrimination index (DI) of neural responses between familiar and novel contexts (Fig. 2D). We found that a significant proportion of neurons discriminated between contexts in both subregions, though a greater proportion did in dCA3 compared to pCA3 (Fig. 2E, F). Using a linear classifier with matched sample sizes, we also found that the information encoded in population activity in dCA3 discriminated between two different contexts with higher accuracy compared to that in pCA3 (Fig. 2G, H). We also evaluated the decoding performance of randomly chosen subsets of recorded neurons from pCA3 (Supplementary Fig. 4E). This analysis showed a correlation between the number of subsampled neurons and peak decoding accuracy, with the performance of the classifier being similar between all recorded pCA3 neurons and dCA3 neurons (Supplementary Fig. 4F).

**Figure 2.**
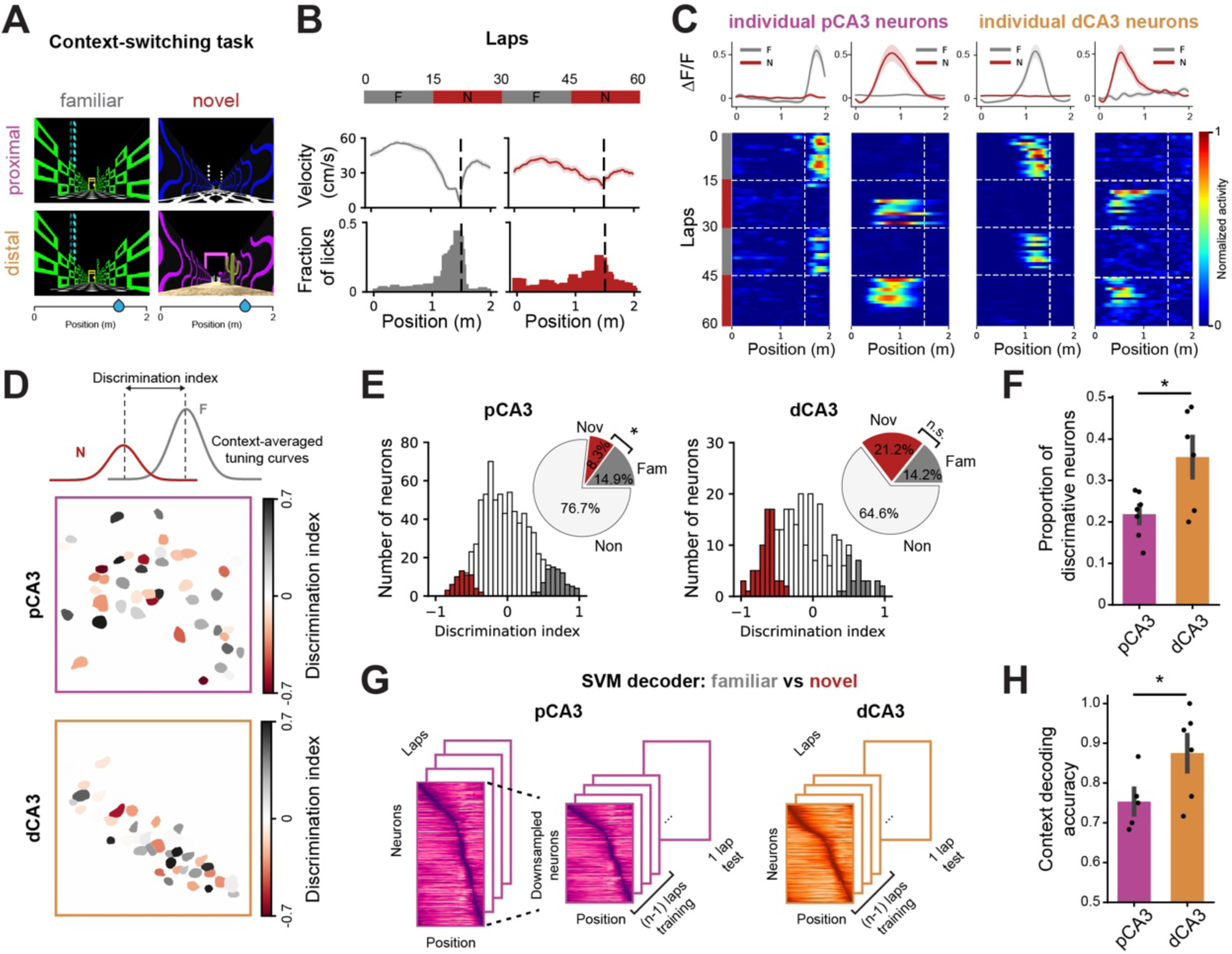
Distal CA3 neurons exhibit more discriminative activity between familiar and novel environments. (A, B) Schematic of the context-switching task (A) and behavioral responses from an example mouse (B). Each session consisted of 60 laps with alternating blocks between familiar (gray) and novel (red) contexts. Both environments were 2m in length with the same location of operant reward. Mice showed different velocity and licking responses in familiar and novel contexts. (C) Mean calcium traces (*top*) and heatmaps of lap-by-lap activity (*bottom*) for example neurons from pCA3 (**B**) and dCA3 (**C**). (D) Calculation of discrimination index (DI) from context-averaged tuning curves (*top*) with the spatial distribution of context-discriminative PNs in pCA3 (*middle*) and dCA3 (*bottom*). (E) Histogram of DI and significant DI values (colored bars, shuffling test, two-sided, p < 0.05). Inset: The fraction of neurons with significant DI in familiar (gray) and novel (red) environments. pCA3 neurons are more significantly tuned for familiar environments (p = 0.035; 7 mice), while dCA3 neurons are not significantly different between environments (p = 0.27; 7 mice). (F) Proportion of neurons with significant DI value (mean ± SEM). dCA3 showed a higher level of context-discriminative neurons compared to pCA3 (p = 0.047; n = 7 mice for both pCA3 and dCA3). (G) Schematic of the support vector machine (SVM) used to decode contextual information. Separate linear decoders were trained for subsampled pCA3 and dCA3 place matrices. (H) SVM linear decoder performance on context discrimination (mean ± SEM). The peak decoding accuracies of dCA3 PNs in discriminating F-N contexts were higher compared to pCA3 PNs (0.61 ± 0.050 for pCA3, 0.85 ± 0.046 for dCA3; p = 0.0011; n = 7 mice for both pCA3 and dCA3). Mann-Whitney U tests were used to determine statistical significance unless otherwise stated. Colors are matched as follows: purple for pCA3, orange for dCA3, gray for familiar context, and red for novel context. n.s. = non-significant, *p < 0.05, **p < 0.01, ***p < 0.001.

In addition to contextual representations, we analyzed place cell dynamics within and between environments. While both pCA3 and dCA3 neurons exhibited spatially tuned responses across context blocks (Fig. 3G, H, Supplementary Fig. 5A), pCA3 place cells showed a higher level of correlation between familiar and novel environments (Fig. 3H). In dCA3, F-N similarity increased during the second switch compared to the first switch, whereas no significant change was observed in pCA3 (Supplementary Fig. 5B, C).

**Figure 3.**
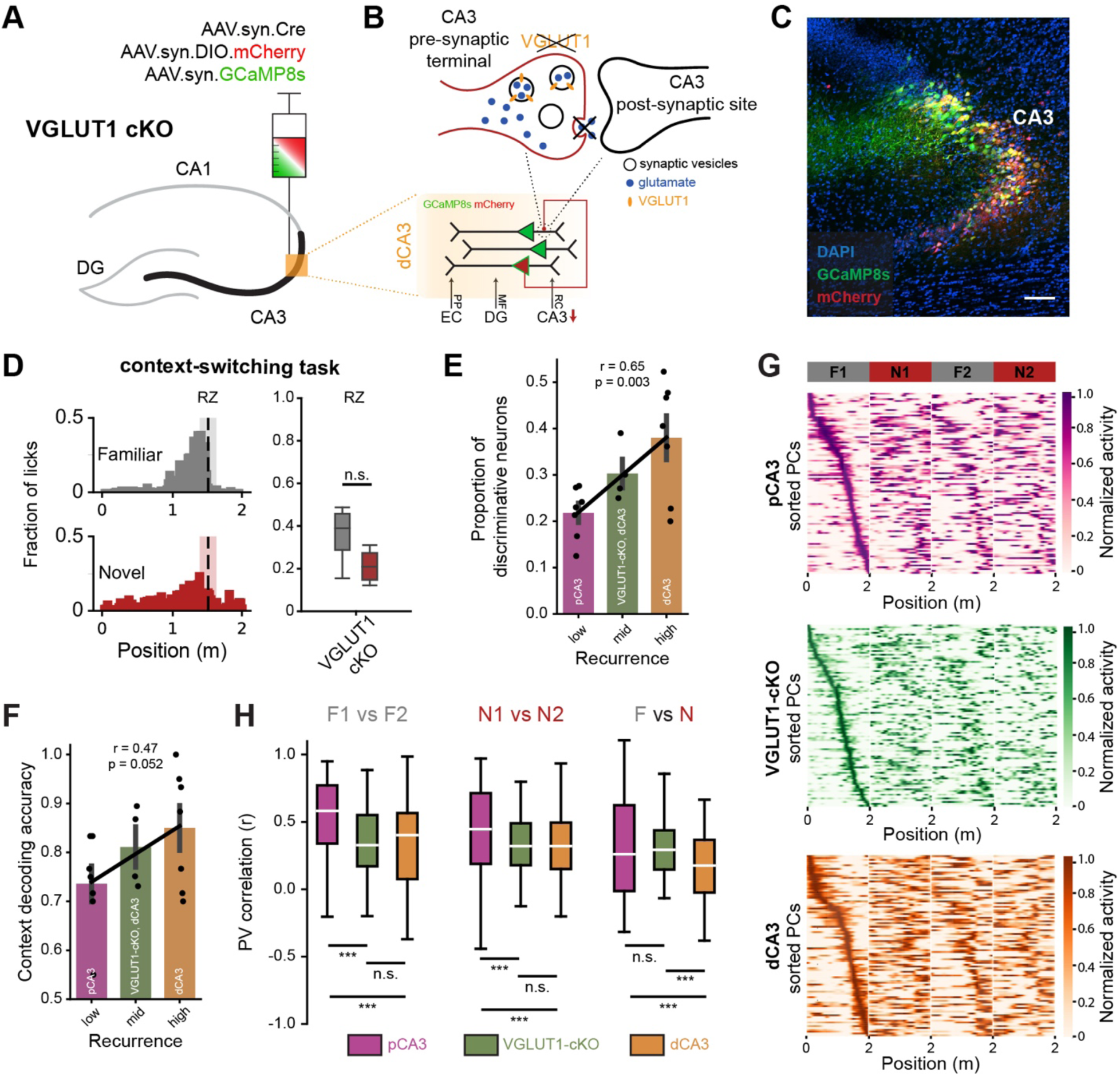
Disrupting glutamatergic synaptic transmission at recurrent collaterals in CA3 alters context-dependent spatial coding. (A) Experimental strategy to disrupt glutamatergic synaptic transmission at CA3 recurrent collaterals using a VGLUT1 conditional knockout (cKO) model. AAV vectors expressing Cre recombinase, DIO-mCherry, and GCaMP8s were injected unilaterally into dCA3 to disrupt and monitor neural activity in the dCA3 region. (B) Schematics illustrating how VGLUT1 deletion affects presynaptic glutamate release onto CA3 PNs. (C) Histological validation of viral expression in dCA3 showing DAPI (blue), GCaMP8s (green), and Cre-dependent mCherry (red). (D) Licking behavior of VGLUT1-cKO mice during the context-switching task, as described in Figure 2A. Left, spatial distribution of licks in familiar and novel contexts; Right, quantification of licking near the reward zone showing no significant difference between contexts (E) Proportion of neurons with significant DI value (mean ± SEM) across recurrence levels (low and high corresponding to pCA3 and dCA3 from Figure 2F; 4 mice for VGLUT1-cKO dCA3; linear regression: r = 0.65, p = 0.003). (F) Context decoding accuracy (mean ± SEM) across recurrence levels (low and high corresponding to pCA3 and dCA3 from Figure 2F; 4 mice for VGLUT1-cKO dCA3; linear regression: r = 0.47, p = 0.052) (G) Heatmaps of spatial tuning curves for pCA3 (top), VGLUT1-cKO (middle), and dCA3 (bottom) PNs during context switching. Rows represent normalized responses sorted by the position of peak activity during the F1 block. (H) PV correlation of spatial tuning curves within contexts (F1 vs F2, N1 vs N2) and between contexts (F vs N). pCA3 showed higher PV correlations within contexts (F1 vs F2 or N1 vs N2). VGLUT1-cKO mice exhibited significantly higher PV correlation between familiar and novel environments compared to dCA3, suggesting disrupted context-specific spatial representations Kruskal–Wallis H tests, F1 vs F2: H = 165.979, p = 9.08×10^-37^; N1 vs N2: H = 55.134, p = 1.07×10^-12^; F vs N: H = 79.937, p = 4.38×10^-18^; Dunn’s post hoc tests, F1 vs F2: p = 2.43×10^-23^ (pCA3 vs VGLUT1-cKO) and 1.80×10^-30^ (pCA3 vs dCA3); N1 vs N2: p = 2.04×10^-7^ (pCA3 vs VGLUT1-cKO) and 1.41×10^-11^ (pCA3 vs dCA3); F vs N: p = 1.20×10^-11^ (pCA3 vs dCA3) and 2.30×10^-15^ (VGLUT1-cKO vs dCA3); n = 7 mice for pCA3 and control dCA3; n = 4 mice for VGLUT1-cKO). Colors: purple, pCA3; green, VGLUT1-cKO; orange, dCA3; gray, familiar; red, novel. n.s., non-significant; *p < 0.05, **p < 0.01, ***p < 0.001.

Taken together, these findings suggest subregion-specific complementary roles for encoding dynamic environments, with pCA3 producing a general template for the context and dCA3 providing more specific responses. Together, these populations might work in concert to balance flexibility and stability for encoding contextual information.

### Characterization of functional heterogeneity along the deep-superficial axis

In addition to the proximodistal axis, previous studies have reported heterogeneity in genetic, morphological, and physiological properties along the deep–superficial axis of hippocampal CA3^15,33^. Whether and how computational properties vary along this axis remains unclear. To address this, we identified superficial and deep CA3 sublayers within our dCA3 imaging fields (see Methods; Supplementary Fig. 6A, B) and compared their coding properties. We found that superficial CA3 neurons exhibited a higher proportion of persistent place cells across days (Supplementary Fig. 6C), whereas tuning stability and context discrimination were similar between sublayers (Supplementary Fig. 6D, E). Given that neuronal subtypes enriched in superficial layers exhibit stronger recurrent connectivity^34,35^, this also aligns with our earlier result showing dCA3 neurons display higher persistence over time (Fig. 1G).

### Disrupting excitatory recurrence alters subregion-specific coding in CA3

Our functional imaging approach revealed dynamic and heterogeneous neural representations along the proximodistal axis of CA3 in relation to both time and context. How two subnetworks of the same hippocampal area implement opposing coding strategies remains an unanswered question. We hypothesized that the observed heterogeneity arises from differences in excitatory recurrent connectivity between the two CA3 subregions.

To test this hypothesis, we selectively reduced glutamatergic transmission in CA3 using a VGLUT1 cKO strategy^27^ and investigated how neural computations change during spatial navigation. By locally co-injecting Cre, Cre-dependent mCherry, and GCaMP8s into dCA3 region of VGLUT1-cKO mice, we selectively reduced VGLUT1-mediated presynaptic glutamate release from recurrent CA3 collaterals while imaging the same neuronal population (Fig. 3A-C). Using immunohistochemistry, we examined the expression of presynaptic glutamatergic markers in dCA3 regions and observed a significant reduction of VGLUT1 expression in the ipsilateral side compared with contralateral control (Supplementary Fig. 7A-C). Despite this substantial change in VGLUT1-labeled fibers, overall neuronal activity levels and place coding property were preserved (Supplementary Fig. 7D, E).

To further assess how reduced recurrence affects contextual processing, we recorded neural activity during a context-switching task (Fig. 3D). Following VGLUT1 cKO, the proportion of context-discriminative neurons was markedly reduced, shifting toward values observed in pCA3 (Fig. 3E). Consistently, SVM decoding accuracy also interpolated between pCA3 and dCA3 (Fig. 3F). Finally, analysis of spatial correlations within and between environments (Fig. 3G, H) revealed that disrupting recurrent excitation significantly increased representational similarity between familiar and novel contexts, biasing coding toward pCA3.

Together, these results provide direct experimental evidence that recurrent excitatory connectivity causally drives the emergence of subregion-specific coding across the CA3 transverse axis.

### Recurrence level interpolates opposing coding strategies

Our causal perturbation experiments demonstrate that recurrent excitation is a key determinant of subregion-specific coding in CA3. To further test whether recurrence level alone is sufficient to shape neural representations and to interpolate between opposing coding strategies, we employed multi-level computational modeling.

First, we constructed two distinct recurrent neural network (RNN) models of CA3: a sparsely connected RNN and a densely connected RNN (Fig. 4A). The differences in recurrent connectivity of the RNNs were chosen based on earlier studies that reported recurrence level heterogeneity across the proximodistal axis of CA3^13,14^ (Supplementary Fig. 8A).

**Figure 4.**
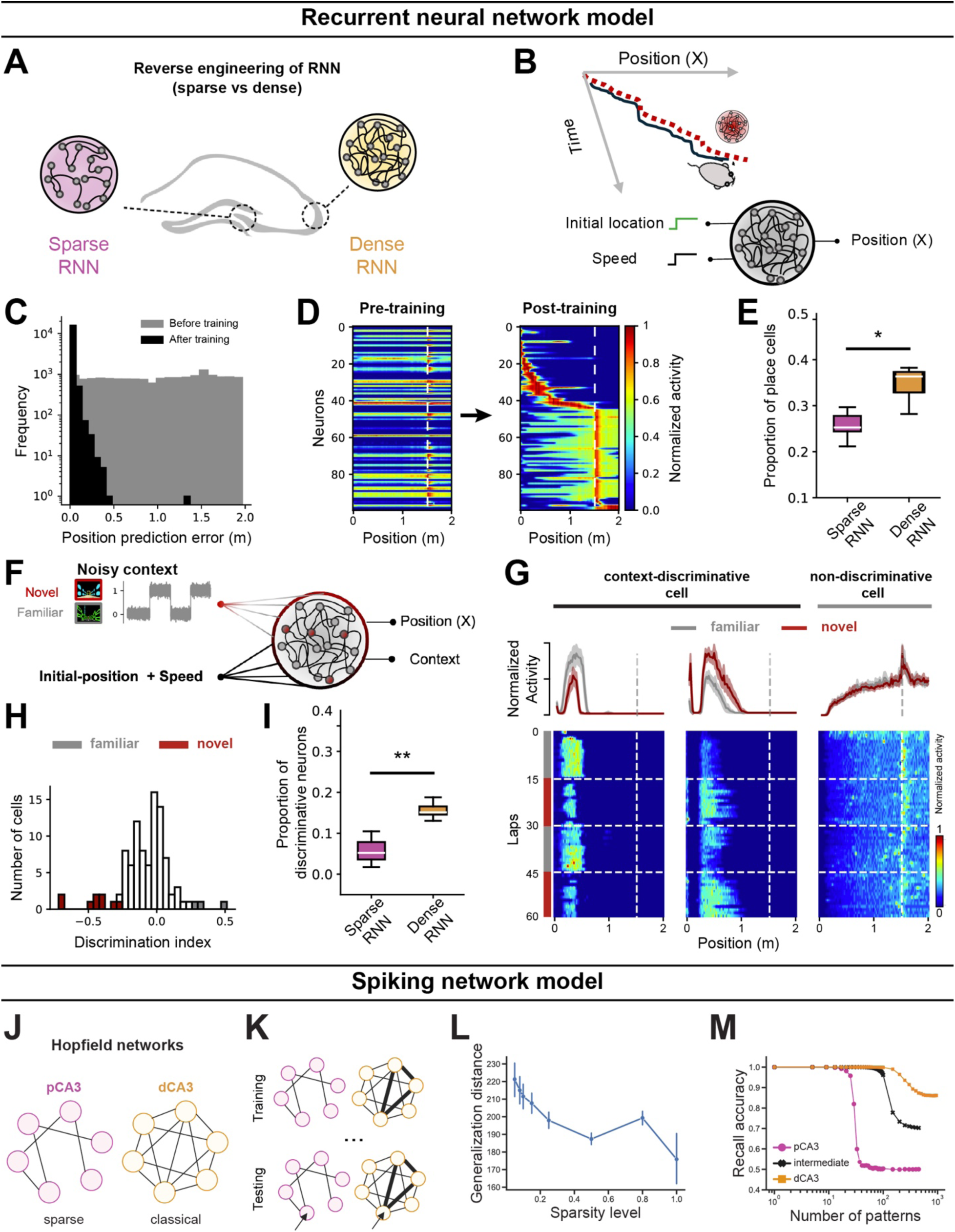
Distinct spatial and memory properties in sparse and dense neural networks. (A) Schematic representation of modeling proximal and distal CA3 with sparse and dense RNNs, respectively. (B) Training of RNNs for a linear track navigation task using inputs of instantaneous speed and lap initiation to mimic mouse trajectory. Network connectivity and activation functions are updated during training, and post-training, activation patterns are analyzed for reverse engineering. (C) Decoding accuracy of the RNNs before (gray) and after (black) training, shown by the position prediction error. (D) Tuning curve evolution of RNNs with 100 cells pre- and post-training, sorted by activity levels. The tuning curves cover the spatial scale of the virtual arena. (E) Proportion of spatial-tuned artificial cells in RNNs with sparse versus dense recurrence. (F) Training of RNNs for context-switching tasks, where task-relevant inputs are fed to RNNs. (G) Examples of artificial cell activation post-training in the context-switching task. The left shows two cells with context modulation (familiar vs. novel), while the right shows a spatially tuned cell without context-dependent activity. (H) Histogram of DI and significant DI values (colored bars, shuffling test, two-sided, p<0.05) for artificial cells. (I) Dependence of context-discriminative cells on recurrence level. Box plots represent the fraction of context-tuned cells for RNNs with sparse versus dense recurrence. (J) Sparse (*left*) and classical (*right*) Hopfield model of associative memory, corresponding to areas pCA3 and dCA3, respectively. (K) Schematic of training (storage) and testing (recall). During storage, binary patterns are sequentially presented to the network and pairwise synaptic weights are updated according to a Hebbian plasticity rule. During retrieval, corrupted patterns are presented to the network, whose dynamics recover the closest previously stored pattern. (L) Sparse networks code more stably than dense networks. 1000-neuron networks were trained to their capacity while still achieving 100% accuracy. Increasingly different patterns were then presented to each network (generalization distance=number of corrupted bits). Sparser networks were able to more accurately recover the underlying memory from patterns with a larger distance to the original, compared to the dense network. This was a trend observed across sparsity levels. (M) Dense network (dCA3) has greater discrimination capacity than the sparse (10% connectivity) network (pCA3). In repeated trials, N patterns are stored in each 1000-neuron network and then recalled (y axis: mean bit-recall accuracy; chance level 50%). Both networks achieve 100% accuracy for small N with accuracy decreasing as N increases, but the number of patterns the sparse network is able to discriminate is much smaller than the dense network. Colors are matched as follows: purple for pCA3, orange for dCA3, gray for familiar context, and red for novel context. n.s. = non-significant, *p < 0.05, **p < 0.01, ***p < 0.001.

Informed by biological evidence showing projections from speed-selective cells in the entorhinal cortex to the hippocampus^36^, we designed our RNN models to receive inputs of speed and the initial location of the animal in a virtual arena (Fig. 4B). The number of training iterations was chosen to ensure both the sparse and dense RNN reached a plateau in their loss functions, ensuring a fair comparison of their performance (Fig. 4C, Supplementary Fig. 8B). Post-training analysis showed that the denser RNN model exhibited a higher proportion of place cells compared to the sparser model, consistent with our experimental observations (Fig. 4D, E, Supplementary Fig. 8C, Supplementary Fig. 1C).

To further explore the adaptability of place cells in the RNNs to new environments, we simultaneously provided a visual context signal and the instantaneous speed of the animal as the inputs to both network models and trained the networks to identify both location and context (Fig. 4F). Given that animals utilize visual cues to identify the context of the VR track^32^, we added noise to the context signal, representing perceptual context evidence from the visual system, to the hippocampal network. We maintained the noise level high enough to avoid a trivial scenario for the network and accordingly reinforced the engagement of a higher proportion of cells in context discrimination. As a result, the RNN model successfully learned the context-switching task with low error rates (Supplementary Fig. 8D); analysis of the hidden layer representations revealed the presence of multiple types of conjunctive and selective cells for context and place (Fig. 4G, H). A greater number of neurons in the dense RNN discriminated between familiar and novel contexts (Fig. 4I), consistent with *in vivo* recording data that showed higher context selectivity in dCA3 (Fig. 2F, H).

Finally, we investigated the effect of varying connectivity on memory generalization and discrimination using a highly abstracted network model of CA3 subregions based on the Hopfield network (Fig. 4J). Hopfield networks are widely used as models of content-addressable memory with powerful theoretical properties. We used Hopfield networks at different sparsity levels, trained with the classic Hebbian rule (Fig. 4K), to model the ability of a CA3 subnetwork to store and retrieve memories^37-39^. We find differences between memory performance in sparsely and densely connected Hopfield networks that mirror differences between pCA3 and dCA3. The dense network exhibited higher memory capacity, i.e., the ability to accurately discriminate a greater number of patterns compared to the sparse network (Fig. 4L). However, the sparse network was better at recovering patterns from significant distortions (Fig. 4M), suggesting fewer, deeper wells in its energy landscape implementing a more stable code. Finally, we find that, consistent with our experimental findings, these abilities interpolate: intermediate levels of connectivity give rise to intermediate levels of discrimination and coding stability (Fig. 4M; black line).

### Dimensionality of neural manifolds altered by recurrence

Theoretical work has established a direct link between recurrency level and representational geometry of neurons, suggesting higher connectivity between neurons increases the mutual activity of cell populations^40^, thereby constraining their representational geometry in population space (Fig. 5A).

**Figure 5.**
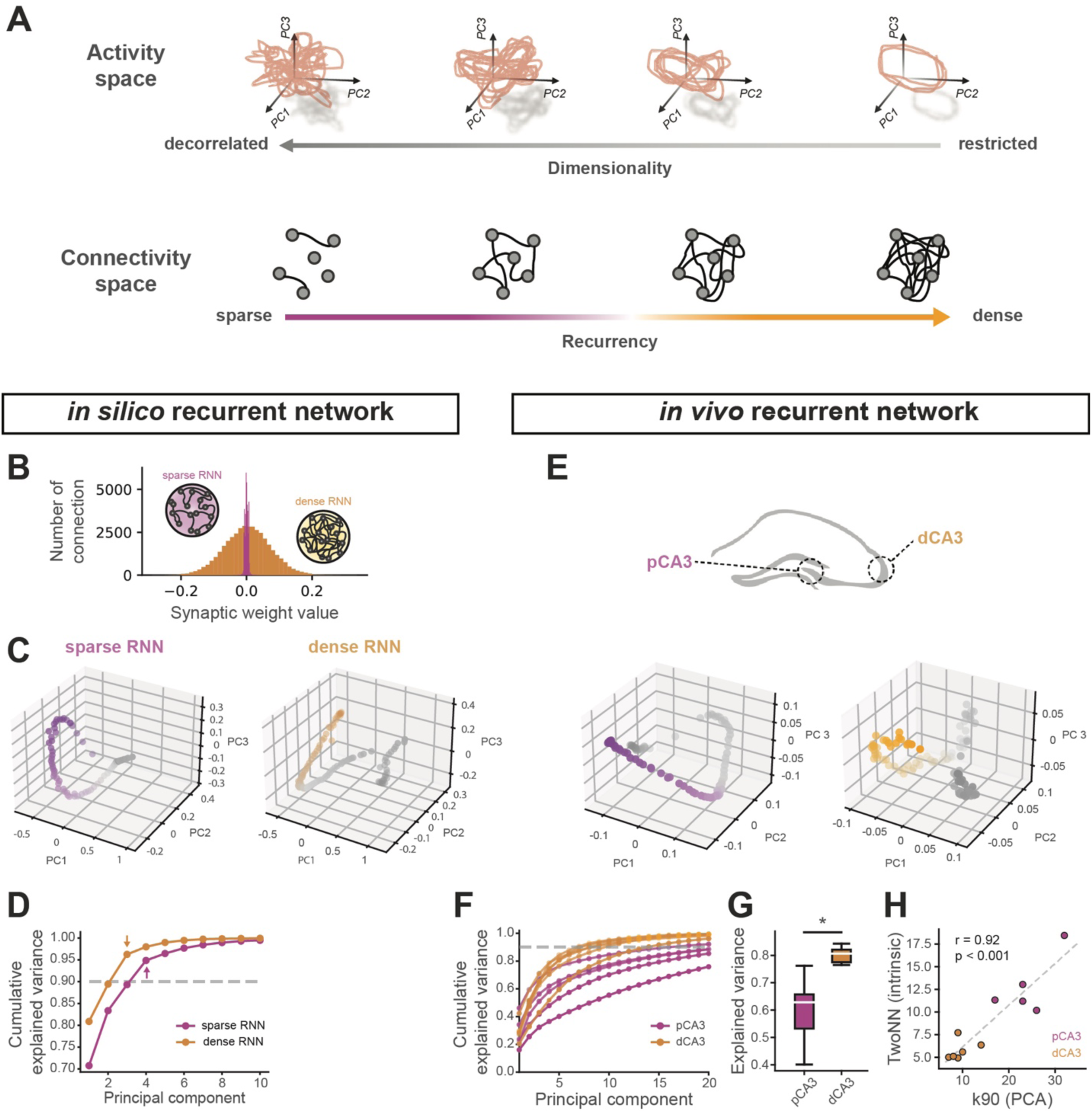
Neural manifold dimensionality in *in vivo* and *in silico* recurrent network. (A) Illustration of a direct link between recurrency and neuronal manifold dimensionality. (B) Distribution of recurrent weights in sparse and dense RNN models. (C) Principal Component Analysis (PCA) of population activities from simulated sparse (*left*) and dense (*right*) RNNs, with projections onto the top three principal components shown. Each dot represents the population activity at a single time point, projected onto PC1–PC3; trajectories color gradient follow progression in task (position). (D) Cumulative explained variance as a function of the number of principal components extracted from PCA with *in silico* simulated datasets. Dashed lines indicate the principal component at which 90% cumulative explained variance is reached. (E) PCA of population activities in pCA3 and dCA3 during spatial navigation tasks, with projections onto the top three principal components shown. (F) Cumulative explained variance as a function of the number of principal components extracted from PCA with *in vivo* datasets. dCA3 curves rise faster than pCA3, consistent with lower dimensionality. Each line represents a mouse and PCA is performed on the full continuous recording rather than on trial-averaged activity, preserving the full temporal dynamics of neural firing on the session-wise neural manifold. Dashed lines indicate the principal component at which 90% cumulative explained variance is reached. (G) Cumulative variance captured by the first 5 PCs. Higher values indicate more concentrated, lower-dimensional representations. (H) Correlation between k90 and TwoNN scatter. The positive correlation confirms agreement between linear (PCA) and nonlinear (nearest neighbor) dimensionality estimators (linear regression: r = 0.92, p = 6.70×10^-6^). Mann-Whitney U tests were used to determine statistical significance unless otherwise stated. Colors are matched as follows: purple for pCA3 or sparse RNN, orange for dCA3 or dense RNN. n.s. = non-significant, *p < 0.05, **p < 0.01, ***p < 0.001.

We first examined this theoretical intuition in simulations of RNNs during the context-switching task. Our simulations demonstrated that altering connectivity towards sparser networks increases the dimensionality of neural manifolds, as defined by the explained variance of cumulative principal components (Fig. 5B-D). This sparsity–dimensionality relationship was preserved across input-drive regimes, activation nonlinearities, and Dale-constrained simulations (k90 and participation ratio; Supplementary Fig. 9). We conducted a similar analysis on the neural activities recorded during the same task to examine the neural manifolds properties of proximodistal CA3 neurons (Fig. 5E). As predicted, the manifold dimensionality of representations in dCA3 was lower than that in pCA3 (Fig. 5F, G), further supporting the role of recurrent connectivity in structuring population-level neural codes within the CA3 circuit. This conclusion was unchanged when dimensionality was estimated using a nonlinear nearest-neighbor intrinsic-dimension metric on trial-averaged trajectories (Fig. 5H, Supplementary Fig. 12).

Recent studies have increasingly highlighted the role of neural representational geometry in supporting cognitive function by balancing specificity and generalization^41,42^. Although our results demonstrate distinct representational dimensionality in dense and sparse networks, how these geometric properties are implemented in spatial and contextual computations in the hippocampus remains unclear. To address this, we analyzed cross-condition generalization performance (CCGP)^43^ to access the alignment of neural manifolds across contexts and positions. Cross-position context CCGP was higher in dCA3 (Supplementary Fig. 4G), whereas cross-context position CCGP was higher in pCA3 (Supplementary Fig. 5D). These findings suggest that the relationship between representational dimensionality and generalization in the hippocampus is not governed by a single rule, but instead depends on the specific task variables (e.g. position, context) being represented.

Together, these findings underscore the critical role of recurrent connectivity in shaping the distinct functional properties of CA3 subregions from single-cell tuning properties to population-level geometry. Our computational modeling, aligned with experimental data, provides deeper insight into the mechanisms by which CA3 PNs contribute to spatial navigation and context discrimination.

## Discussion

Despite extensive evidence of anatomical, molecular, and physiological heterogeneity in the CA3 along the transverse axis^13-16,44,45^, the relationship between these differences and neural computations performed by CA3 subregions has remained largely unknown. While both stable and flexible dynamics as well as varying degrees of context generalization have been reported in CA3 using *in vivo* imaging^46-49^, these representational features have not been systematically investigated along the proximodistal axis, nor have they been linked to CA3 subregions. In this study, we bridged these gaps by combining *in vivo* two-photon calcium imaging of CA3 subregions with computational modeling at multiple levels of abstraction. Our findings provide significant insights into how distinct hippocampal subcircuits differentially contribute to memory functions^46,50,51^.

We found that dCA3 neurons are highly sensitive to contextual changes as well as tuned to different locations over days. This suggests that they are flexible and adaptive, possibly playing a role in learning and responding to new experiences or changes in the environment. In contrast, pCA3 neurons show more stability in their tuning over time, even though fewer neurons are tuned compared to dCA3. This stability, consistent with a previous report^46^, suggests that they might be involved in encoding persistent or core aspects of the context that remain relevant across different experiences. These neurons may be essential for maintaining a stable representation of the environment, supporting consistent recall of key contextual features or stable memory traces, and could be rooted in how they integrate inputs at the subcellular level^52-54^.

Previous studies have demonstrated distinct computational processes across hippocampal regions, such as pattern separation in the DG and pattern completion in CA3^11,55^, despite recent evidence that CA3 can support both processes^17^. These classic frameworks employed tasks with gradual or partial cue manipulations to probe responses to overlapping inputs. In contrast, our study used discrete, highly distinct contexts (F vs. N), shifting the focus from completion or separation of similar inputs to neural discriminability between distinct environments. Under these conditions, we observe subregion-specific differences in the stability and flexibility of CA3 representations at both single-cell and population levels which persisted across days. Thus, our results do not contradict prior work but instead complement it by revealing how CA3 coding strategies depend on local circuit structure and task demands.

In addition to the proximodistal axis, the hippocampus exhibits prominent radial-layer organization, yet deep–superficial heterogeneity has been primarily characterized in CA1^56,57^. Our findings extend this principle to CA3 by showing that superficial neurons display greater long-term place cell persistence than deep neurons. Considered together with the higher persistence observed in distal CA3, these results highlight the critical role of recurrent connectivity in supporting representational stability across CA3 subcircuits. Although our current approach does not resolve morphologically defined CA3 subtypes^33,34^, future work combining cell-type–specific labeling could directly test whether CA3 pyramidal subtypes exhibit distinct coding dynamics along the anatomical axis *in vivo*.

Consistent with recent reports on the rapid formation of place fields in the hippocampal CA1 and CA3^31,47,48,58^, we also found that both pCA3 and dCA3 PNs exhibit spatially tuned neural responses that rapidly formed when the mice are exposed to novel environments. Future studies will explore the synaptic plasticity rules governing the dynamics of place field formation in CA3 along the transverse axis^59^. We further observe that dCA3 PNs are more significantly tuned to novel environments compared to pCA3 PNs. It is well known that hippocampal circuits are under neuromodulatory control related to the behavioral state of animals^60-64^. Correspondingly, the CA3 subregions may receive different levels of catecholaminergic input, such as dopamine and norepinephrine from the locus coeruleus, which facilitates memory formation in a new context^65-67^.

Using targeted genetic manipulation of excitatory connectivity, we provide direct causal evidence that excitatory recurrence contributes to the distinct coding strategies of proximal and distal CA3. By selectively reducing glutamatergic transmission in dCA3, we found that its coding properties shifted toward those typically observed in pCA3, with a reduced proportion of context-discriminative neurons. Importantly, this manipulation provides *in vivo* evidence for the predictions of our modeling framework: the shift from dynamic to stable codes when recurrence is knocked down closely parallels predictions from both our RNN and Hopfield network simulations. Although hippocampal recurrent connectivity is substantially sparser than that of cortical circuits^34,38,68,69^, our manipulation experiments show that small changes within a sparse network can nonetheless induce pronounced effects on network dynamics. Consistent with this, our modeling approaches further demonstrate that sparsity emerges as a sensitive control parameter that strongly shapes manifold dimensionality and recall accuracy (Supplementary Fig. 9, 11). Importantly, this does not imply that other circuit parameters are negligible. In the modeling framework, the sparsity-dependent regime shift was preserved across systematic variations in input drive, neuronal nonlinearities, and Dale-constrained E/I structure (Supplementary Fig. 9). Together, the combination of targeted circuit manipulation, population imaging, and modeling provides convergent evidence that heterogeneity in recurrent connectivity is a key determinant of functional specialization within CA3.

It is striking that two subregions of the same hippocampal area, separated by less than a millimeter in the mouse brain and (notwithstanding the important differences we have highlighted) broadly similar in their anatomy, physiology, and development, can implement seemingly opposing computational functions. Our RNN modeling results, in which spatial and contextual representations by artificial cells corresponded closely to the empirical data in both single-cell and population level, underscore the critical role of local connectivity patterns in shaping the functional properties of neural networks. Furthermore, simulation results with dense, intermediate, and sparse Hopfield networks imply a smooth gradient between dynamic and stable codes based on the level of recurrence in a subnetwork. Taken together, these findings emphasize the specific importance of recurrent connectivity in determining the computational features of neural circuits, supporting the idea that small variations in structural connectivity can lead to substantial differences in network function and behavior. Crucially, this amplification of small structural differences is not uniform across connectivity regimes: simulations spanning the full sparsity landscape show that networks in sparse regimes (comparable to hippocampal regions) exhibit significantly greater sensitivity to small connectivity perturbations than dense networks, with even a few-percent change in sparsity producing measurable shifts in manifold dimensionality (Supplementary Fig. 13).

Our findings show that recurrent connectivity shapes the dimensionality of neural manifolds in CA3, with sparser networks yielding higher-dimensional manifolds and denser networks producing lower-dimensional ones. While an increasing body of literature associates dimensionality in neural spaces with flexibility and generalization^41,70-72,^ dimensionality alone does not fully account for a network’s computational capacity. The geometry of neural representations—encompassing factors like curvature, separation, and topology—plays a critical role in how information is processed^73-77^. To fully understand flexibility and generalization, it is necessary to study the precise geometry of these representations. Previous studies have shown divergent roles in task performance are observed in networks with varying neural trajectory geometries^78,79^.

Recent theoretical studies have explored how structural connectivity within networks affects their representational space^80,81^, in addition to the computational works examining this structure-to-function interaction in low rank regimes^82,83^. However, whether biological networks are inherently low-rank remains under debate. In our approach, we focus on binary (present-or-absent) connectivity, providing a clearer structural interpretation of how network architecture shapes representational dimensionality, complementing earlier studies that link structural biases to neuronal variability^84,85^. Theory predicts that low-dimensional activity remains possible in sparse networks through fine-tuning of synaptic weights^86^. Accordingly, the direct measurement of synaptic weights *in vivo* will be an important direction for future experimental work to dissect the synaptic basis for computational capacity.

CA3 PNs are well known to exhibit burst firing *in vivo^33,87,88^*. Despite the improved kinetics of GCaMP8s^89^, our approach does not fully resolve the temporal structure of bursts. Given the strong impact of burst activity on synaptic plasticity and network dynamics, future studies combining optical recordings with simultaneous electrophysiology or high-speed voltage imaging approaches^90^ will be critical for dissecting the specific role of burst firing in CA3 representations.

Because CA3 is a high-dimensional, heterogeneous circuit, mechanistic modeling necessarily requires abstraction: simplified networks are used not as biophysically faithful replicas, but as controlled testbeds for counterfactual reasoning about which circuit variables are sufficient to generate qualitative coding regimes. This approach is widely used in neuroscience, including work showing how abstract recurrent networks can capture key dynamical motifs and computations despite missing biological detail^91,92^, and broader perspectives emphasizing the value of goal/task-driven artificial networks as mechanistic models at the level of representations and population dynamics^93^. Accordingly, we interpret the RNN/Hopfield results as a mechanistic sufficiency test for recurrent sparsity as a contributing axis of population geometry and coding regime, rather than as a quantitative fit to CA3 microcircuit statistics.

In addition to recurrent local connectivity, subregion-specific differences in inputs to CA3 can contribute to the observed differences in context-related dynamics between pCA3 and dCA3. Recent reports have highlighted a generalized neural representation by dentate gyrus granule cells, one of the main input sources to CA3^46^, which may contribute to the more generalized contextual and spatial dynamics we observe in pCA3 PNs compared to dCA3 PNs. Similarly, subregion-specific differences in inputs from the entorhinal cortex, which is also implicated in context generalization and discrimination in the CA3^94,95^, may also shape context processing in CA3. Finally, the role of inhibition and subtype-specific inhibitory control of feature representations and their stability remain unknown^96-98^. Future studies should investigate how external inputs modulate the cognitive flexibility of different subregions within CA3. Understanding local circuit^99^ and long-range interactions will provide deeper insights into how the hippocampus supports complex behaviors, integrating contextual and spatial information to facilitate adaptive navigation and memory.

## Supplementary Figures

**Supplementary Figure 1.**
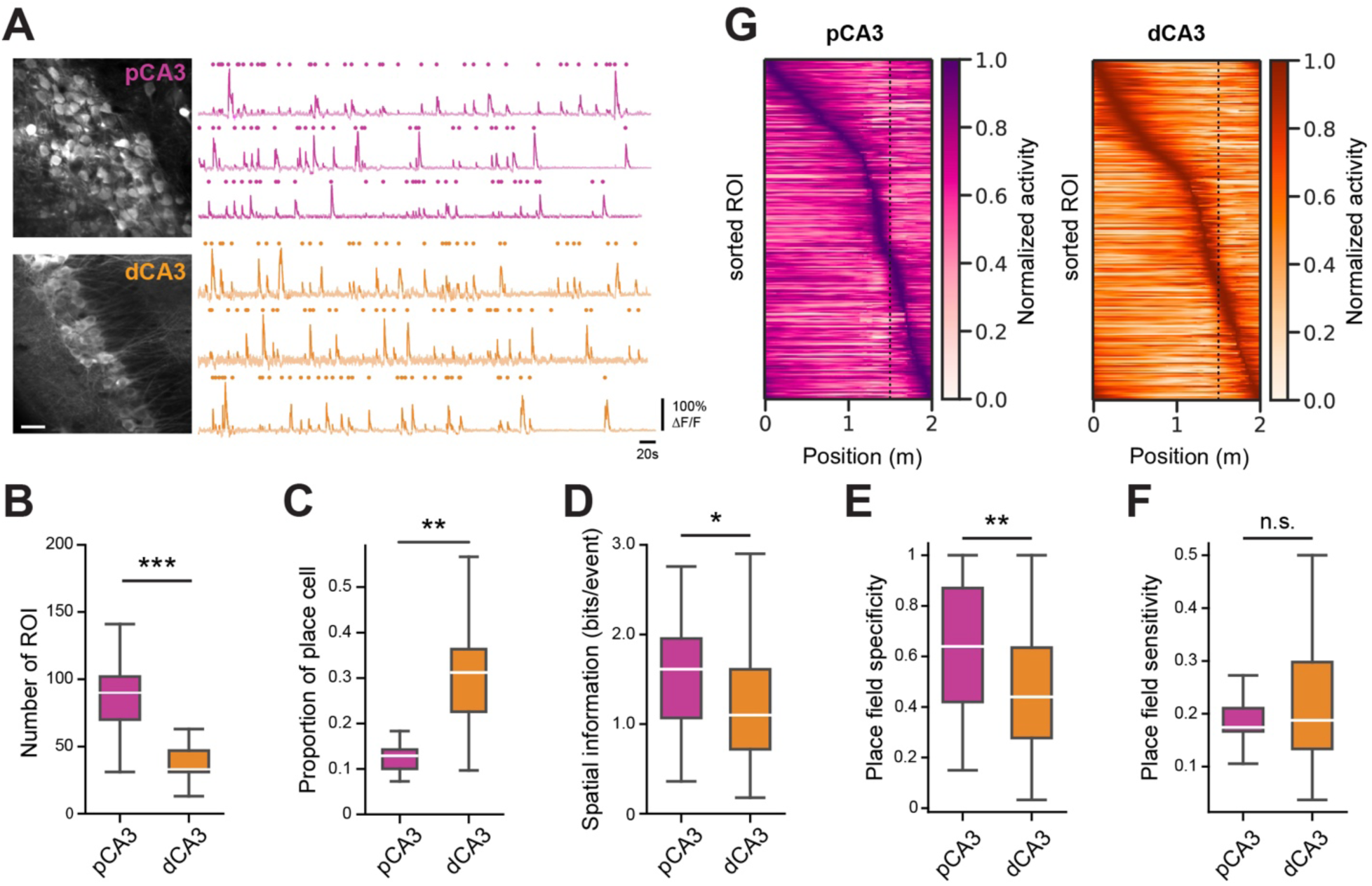
(Related to Figure 1) Spatial tuning of proximal and distal CA3 during spatial navigation. (A) Representative time average image (*left*) and ΔF/F traces (*right*) of proximal (*top*) and distal (*bottom*) CA3 neurons. Transient events in bold and start times indicated by circles above traces. (B) Average number of detected ROI in proximal and distal CA3 (p = 3.25×10^-5^). (C) Proportion of place cells in each CA3 subregion (mean ± SEM; p = 0.0025; n = 5 and 7 mice for pCA3 and dCA3, respectively). (D) Spatial information (bits/event) of place cells in each subregion (p = 0.037) (E) Place field specificity in each subregion (p = 0.0076). (F) Place field sensitivity in each subregion (p = 0.58). (G) Heatmaps of spatial tuning curves for pCA3 (*left*) and dCA3 (*right*) cells. Rows show average and normalized responses for all cells, sorted by the position of the peak activity. Mann-Whitney U tests were used to determine statistical significance unless otherwise stated. Colors are matched with purple for pCA3 and orange for dCA3. n.s. = non-significant, *p < 0.05, **p < 0.01, ***p < 0.001.

**Supplementary Figure 2.**
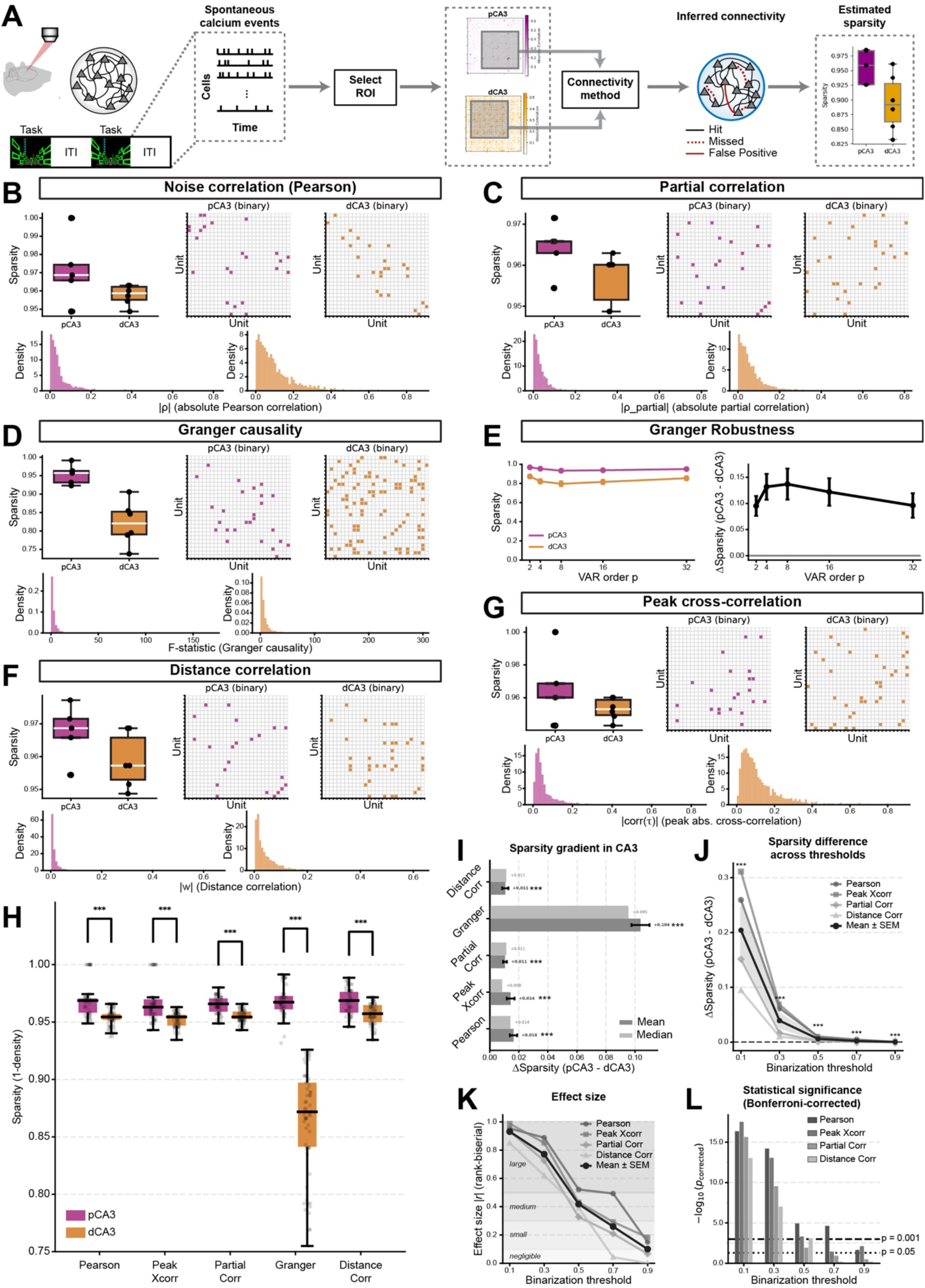
(Related to Figure 1) Heterogeneity of recurrent connectivity level along the proximodistal axis of CA3. (A) Connectivity estimation pipeline. Spontaneous calcium events are recorded from hippocampal CA3 neurons during inter-trial intervals (ITI). ROIs are selected, yielding a neurons × time activity matrix for each session to avoid cell number imbalance across regions. Five connectivity estimation methods are applied independently to infer pairwise functional connectivity. Each method produces a continuous weight matrix, which is binarized using an adaptive threshold (μ + 2σ of off-diagonal absolute weights) or, for Granger causality, a Bonferroni-corrected p-value threshold. Network sparsity is then computed for each session based on the resulting binary adjacency matrix. (B) Pairwise Pearson correlation reveals sparser functional connectivity in pCA3 than dCA3. (*Top*) Box plots of network sparsity for pCA3 and dCA3, computed from absolute Pearson correlation matrices binarized with an adaptive threshold (μ + 2σ of off-diagonal |ρ|). Each dot represents one recording session (n = 5 and 6 for pCA3 and dCA3, respectively). Center and right panels show example binary adjacency matrices from one representative session per area, where colored entries indicate suprathreshold connections. pCA3 networks are visibly sparser. (*Bottom*) Histograms of all off-diagonal absolute Pearson correlation coefficients (|ρ|) pooled across sessions for pCA3 (*left*) and dCA3 (*right*). (C) Partial correlation (Ledoit–Wolf regularization) isolates direct pairwise connections and confirms sparser pCA3 networks. (D) Granger causality detects sparser directed functional connectivity in pCA3. Unlike correlation-based methods, Granger causality captures directed (asymmetric) temporal dependencies: whether past activity of neuron ⅈ predicts future activity of neuron ⅉ beyond ⅉ’s own history. (E) Granger robustness across VAR model orders. *(Left)* Mean ± SEM sparsity for pCA3 and dCA3 as a function of VAR model order p ∈ {2, 4, 8, 16, 32}. At every tested order, pCA3 lies above dCA3. *(Right)* The difference Δ Sparsity (pCA3 − dCA3) ± SEM remains positive across all model orders, demonstrating that the sparser pCA3 connectivity is not an artifact of the chosen temporal lag structure. (F) Distance correlation, a nonlinear dependency measure, confirms sparser pCA3 networks. Even when nonlinear dependencies are captured, pCA3 displays sparser functional coupling than dCA3, ruling out the possibility that the sparsity difference is limited to linear co-activity patterns. (G) Peak cross-correlation across temporal lags yields sparser pCA3 connectivity. Accounting for temporal delays between neuron pairs does not eliminate the pCA3–dCA3 sparsity difference, indicating that the finding is not driven by synchrony at a single time lag. (H) pCA3 is consistently sparser than dCA3 across all five connectivity estimation methods. To increase statistical power, 10 random contiguous windows (50% of each session’s frames) are drawn per session, and connectivity is computed on each window independently for all 5 methods. (I) Sparsity gradient. Horizontal bars show Δ Sparsity (pCA3 − dCA3) for each method (mean difference ± SEM). (J) Threshold Robustness Analysis. The pCA3 > dCA3 sparsity difference is robust to binarization threshold choice. To confirm that the sparsity finding is not driven by a particular threshold, four correlation-based methods (Pearson, peak cross-correlation, partial correlation, distance correlation) are swept across five absolute binarization thresholds (0.1, 0.3, 0.5, 0.7, 0.9). Granger causality is excluded because it uses p-value–based binarization. Each method × threshold combination uses window resampling (N = 10 windows, 50% of frames) on neuron-equalized data. All traces remain above zero at every threshold (Fisher’s method). (K) Effect size. Rank-biserial effect size |r| (derived from Mann–Whitney U) for each method at each threshold. Reference bands indicate standard regimes (negligible: 0–0.1; small: 0.1–0.3; medium: 0.3–0.5; large: >0.5). Effect sizes are consistently medium-to-large across the full threshold range, indicating practically meaningful differences rather than merely statistically significant ones. (L) Statistical significance (Bonferroni-corrected). Grouped bar chart of −log₁₀(p_corrected_) at each threshold, with Bonferroni correction for multiple threshold comparisons. Horizontal reference lines mark p = 0.05 and p = 0.001. Nearly all method × threshold combinations exceed the p = 0.001 significance line. *Sign consistency:* 95% (19/20) of all method × threshold combinations show pCA3 > dCA3 (Δ > 0). Mann-Whitney U tests were used to determine statistical significance unless otherwise stated. Colors are matched as follows: purple for pCA3, orange for dCA3. n.s. = non-significant, *p < 0.05, **p < 0.01, ***p < 0.001.

**Supplementary Figure 3.**
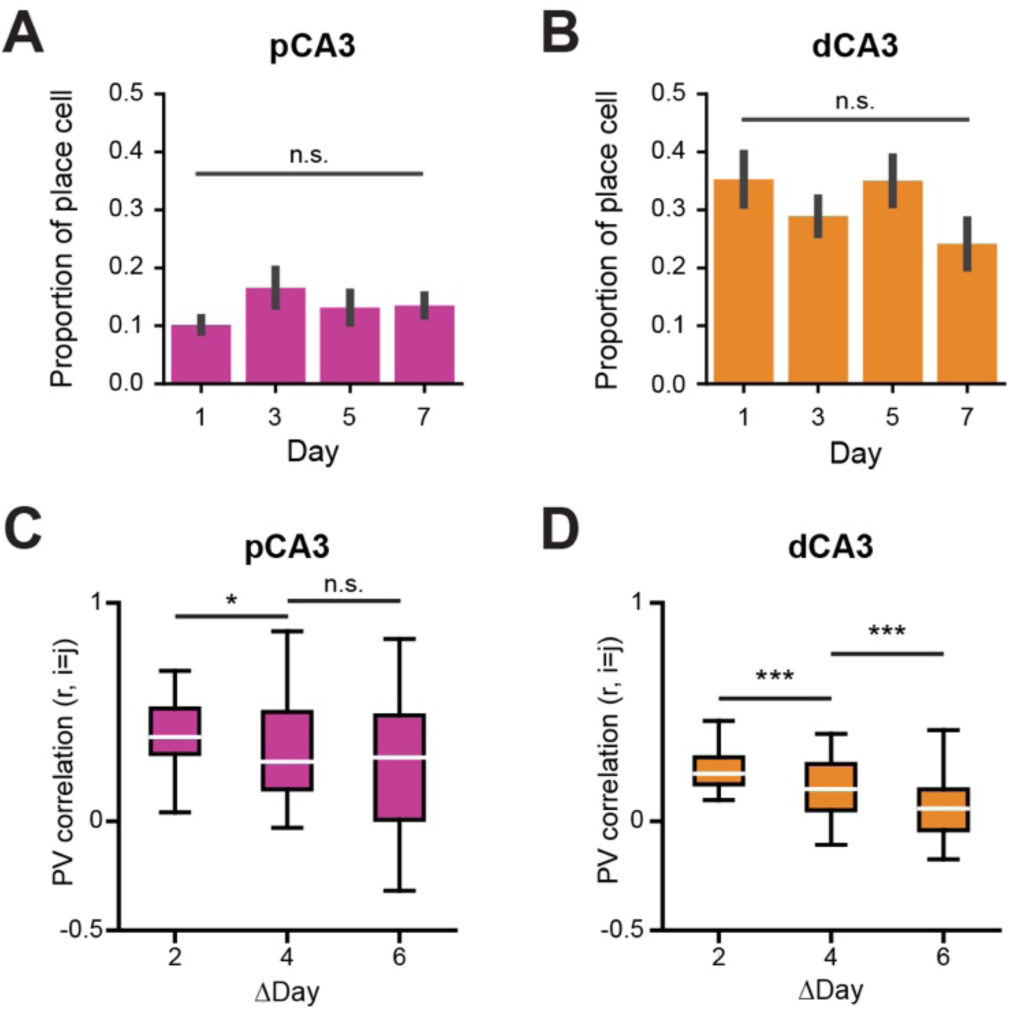
(Related to Figure 1) Long-term stability of place cells in proximodistal CA3. (A, B) Proportion of place cells across days in pCA3 (A) and dCA3 (B). No significant difference was observed across days in either region (Kruskal-Wallis *H*-test, pCA3: H = 2.69, p = 0.91; dCA3: H = 2.70, p = 0.72; post-hoc Dunn’s multiple comparison with Bonferroni correction). (C, D) PV correlation between the same position across days of place cells from pCA3 (C) and dCA3 (D). Correlation values decreased as interval between days increased. (C) p = 0.015 between intervals 2 and 4, p= 0.088 between intervals 4 and 6. (D) p = 6.81×10^-6^ between intervals 2 and 4, p = 0.00014 between intervals 4 and 6. Two-sided unpaired t-tests were used to determine statistical significance. Colors are matched with purple for pCA3 and orange for dCA3. n.s. = non-significant, *p < 0.05, **p < 0.01, ***p < 0.001.

**Supplementary Figure 4.**
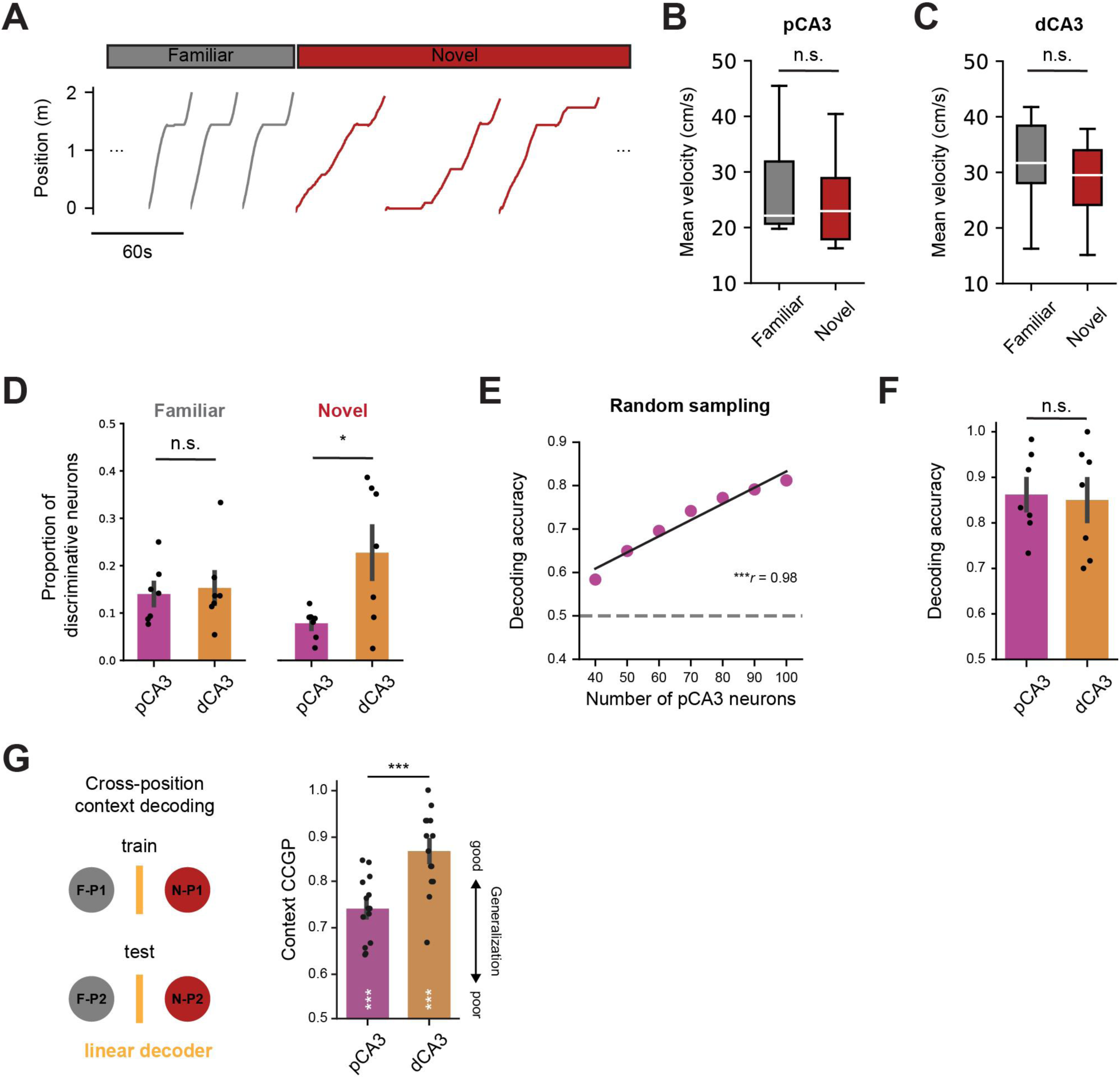
(Related to Figure 2) Context-discriminative neural activity in proximodistal CA3. (A) Position of the animal in the familiar and novel environments from an example session. (B, C) Comparison of mean velocity in the familiar and novel environments for pCA3 (B) (p = 0.53; n = 7mice) and dCA3 (C) (p = 0.32; n = 7mice) recording sessions. (D) Proportion of neurons with significant DI value in familiar and novel environments (7 mice for both pCA3 and dCA3). The proportion of PNs significantly tuned to the familiar environment was similar between pCA3 and dCA3 (0.14 ± 0.23 for pCA3, 0.15 ± 0.03 for dCA3; p = 1.0), but dCA3 showed a higher level of novelty-tuned neural responses compared to pCA3 (0.078 ± 0.012 for pCA3, 0.23 ± 0.055 for dCA3; p = 0.021). (E) Peak decoding accuracy correlated with the number of randomly subsampled pCA3 neurons (Pearson’s *r* = 0.98, p = 0.00015, 7 mice). The black line indicates the regression line. (F) Performance of the linear decoder for context in pCA3 and dCA3 without subsampling. No significant differences were found in peak decoding accuracies between pCA3 and dCA3 (0.86 ± 0.034 for pCA3, 0.85 ± 0.046 for dCA3, p = 0.85, n = 7 mice for both pCA3 and dCA3). (G) Schematic of decoding and CCGP for context (left; see Methods) and CCGP decoding results (right). A linear decoder was trained on neural activity from one spatial bin (F-P1 vs N-P1) and tested on a non-overlapping bin (F-P2 vs N-P2) to assess cross-position generalization of context representation. Context CCGP was significantly higher in dCA3 than in pCA3 (0.74 ± 0.02 for pCA3, 0.87 ± 0.02 for dCA3, p = 0.00085, n = 14 sessions for both pCA3 and dCA3); white asterisks indicate significance relative to the shuffled distribution (p = 9.71×10^-9^ for pCA3; p = 1.08×10^-9^ for dCA3). Mann-Whitney U tests were used to determine statistical significance unless otherwise stated. Colors are matched as follows: purple for pCA3, orange for dCA3, gray for familiar context, and red for novel context. n.s. = non-significant, *p < 0.05, **p < 0.01, ***p < 0.001.

**Supplementary Figure 5.**
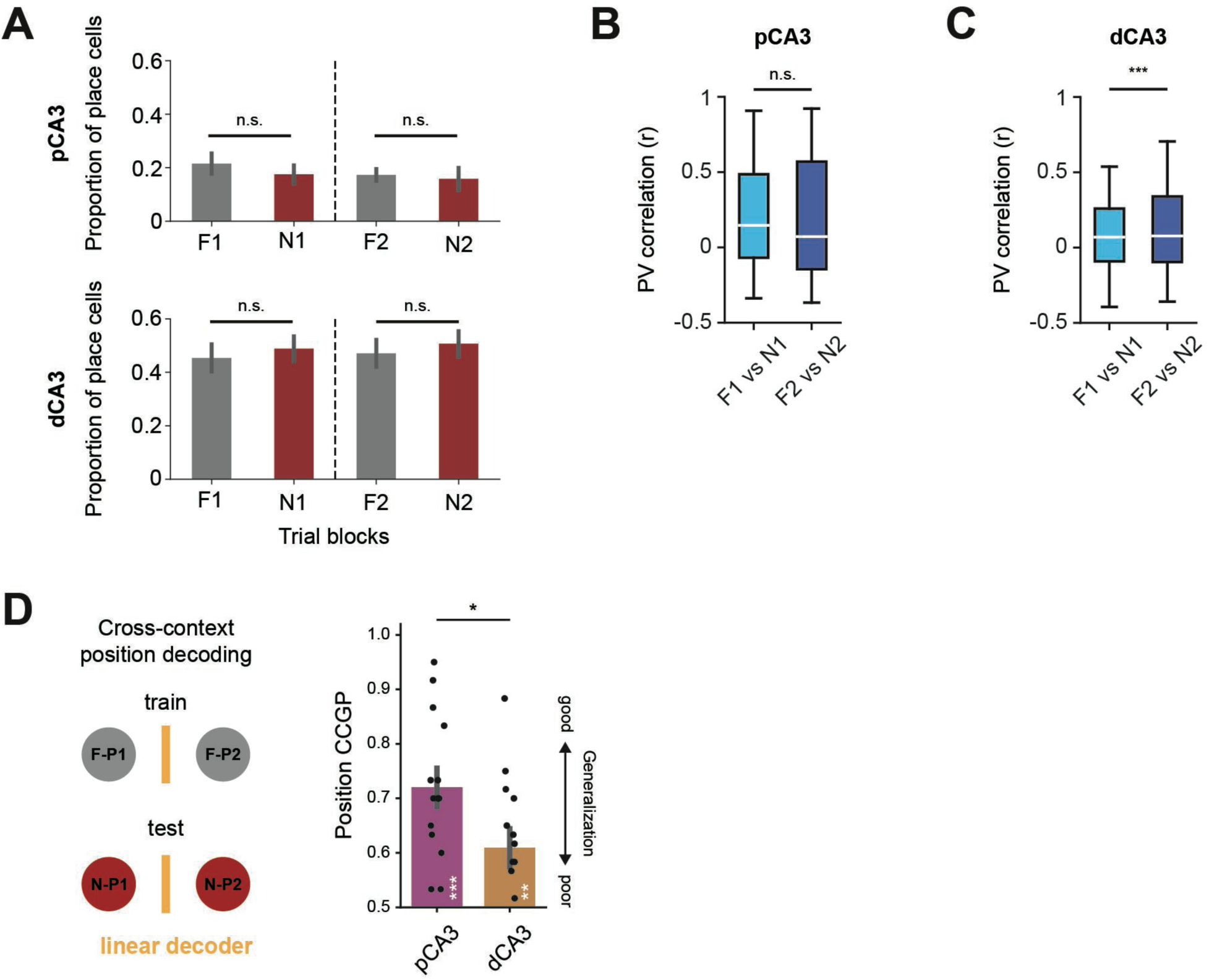
(Related to Figure 3) Place cell responses across blocks in a context-switching task. (A) Proportion of place cells in each trial block in pCA3 (top) and dCA3 (bottom). Familiar and novel blocks showed similar levels of place cells in both the first and second exposures (Kruskal-Wallis H-test, H = 1.43, p = 0.69 for pCA3, H = 0.73, p = 0.87 for dCA3). (B, C) PV correlation between familiar and novel contexts for the first and second blocks. No significant difference was observed in pCA3 between the first and second blocks (p = 0.71), whereas a significant difference was found in dCA3 (p = 0.0003). (D) Schematic of decoding and CCGP for position (left; see Methods) and CCGP decoding results (right). A linear decoder was trained on neural activity distinguishing two spatial locations (P1 and P2) in one context (familiar) and tested on the other context (novel), and vice versa, to assess cross-context generalization of positional representation. Position CCGP was significantly higher in pCA3 than in dCA3 (0.72 ± 0.04 for pCA3, 0.61 ± 0.04 for dCA3, p = 0.04, n = 14 sessions for both pCA3 and dCA3); white asterisks indicate significance relative to the shuffled distribution (p = 2.84×10⁻⁵ for pCA3; p = 0.0076 for dCA3) Colors are matched as follows: purple for pCA3, orange for dCA3, gray for familiar context, and red for novel context. n.s. = non-significant, *p < 0.05, **p < 0.01, ***p < 0.001.

**Supplementary Figure 6.**
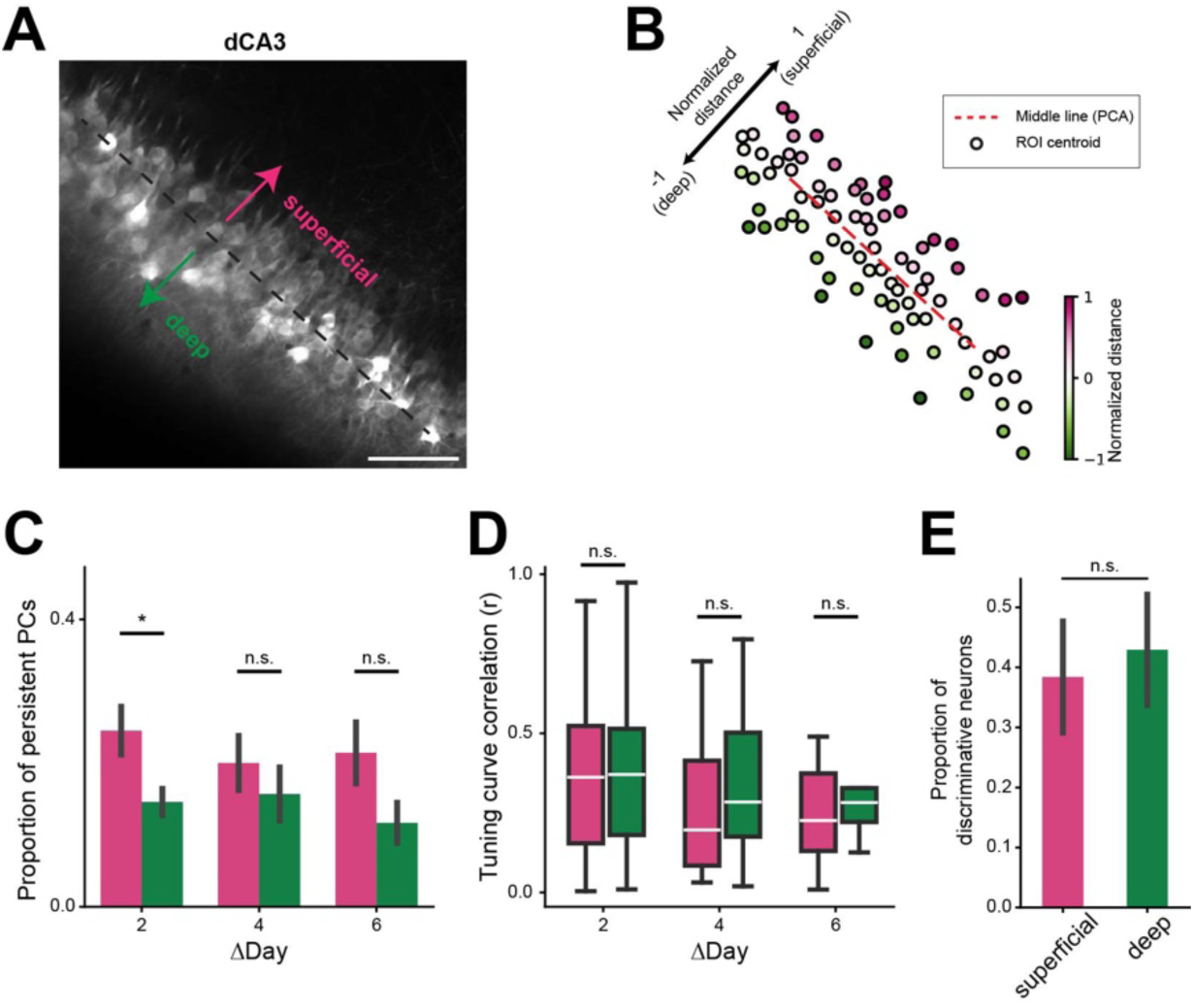
(Related to Figure 1 and 2) Spatial coding properties of CA3 pyramidal neurons along the deep-superficial axis. (A) Example two-photon imaging field of view showing the deep-superficial axis in dorsal CA3. Arrows indicate the superficial (pink) and deep (green) directions. Scale bar: 50 um. (B) Example ROI centroid map from the same dataset shown in (A), color-coded by normalized signed distance from the PC1-derived middle line (see Methods: Deep–superficial cell classification). Positive values correspond to superficial positions (top 10%), negative values to deep positions (bottom 10%). (C) Proportion of persistent place cells across days for superficial (pink) and deep (green) groups (mean ± SEM; p = 0.013, 0.042, 0.080 for ΔDay 2, 4, and 6, respectively). Persistent PCs were defined as neurons classified as place cells in both sessions, as in Figure 1G. (D) Single-cell tuning curve correlation (Pearson’s *r*) across days for persistent PCs in superficial and deep groups (p = 0.86, 0.31, 0.31 for ΔDay 2, 4, and 6, respectively;). (E) Proportion of context-discriminative neurons in superficial and deep groups (mean ± SEM; p = 0.56). Mann–Whitney U tests were used to determine statistical significance unless otherwise stated. Colors are matched as follows: pink for superficial and green for deep. n.s., not significant; *, p < 0.05; **, p < 0.01; ***, p < 0.001.

**Supplementary Figure 7.**
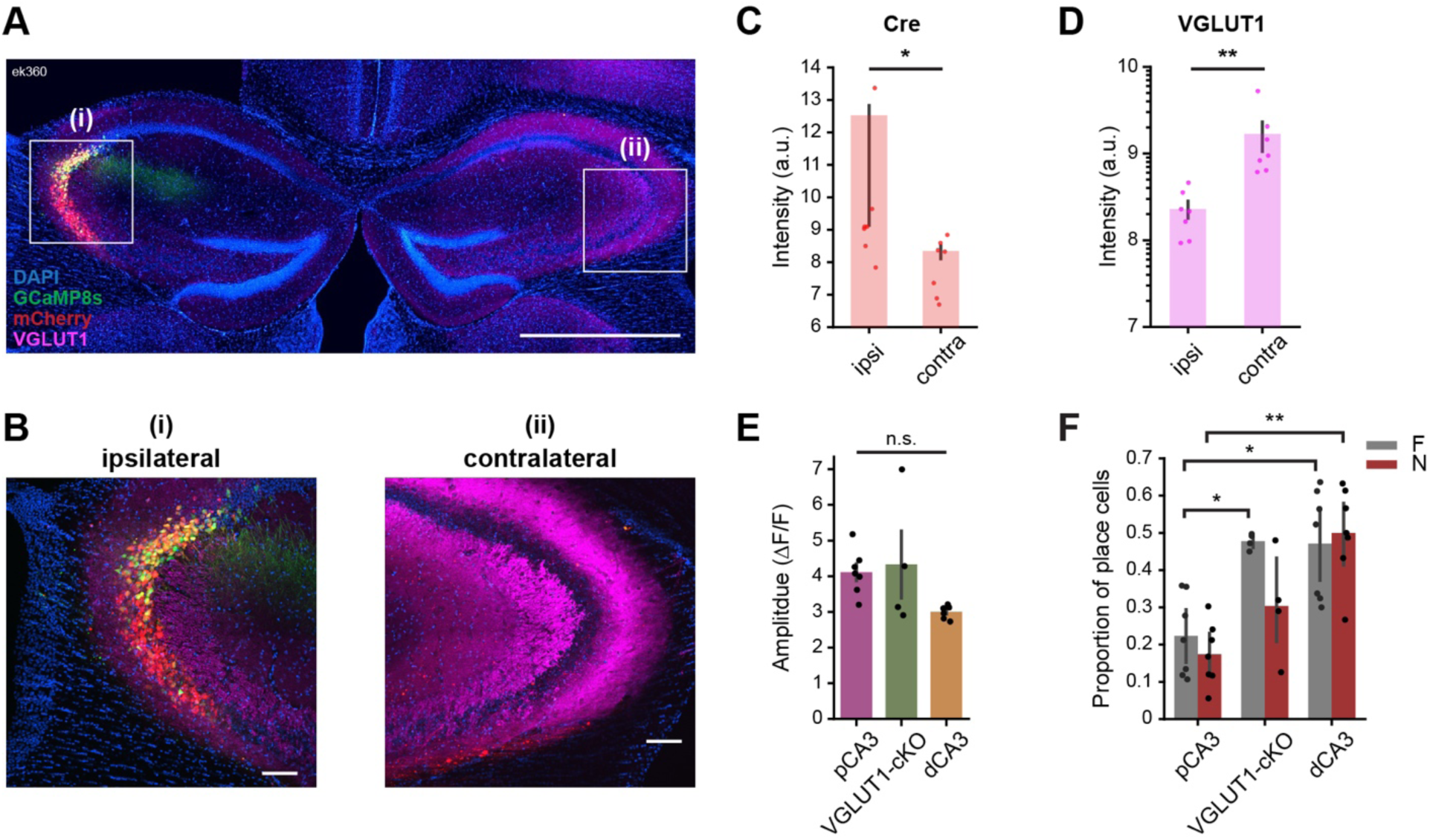
(Related to Figure 3) Validation of VGLUT1-cKO and effects on neural activity. (A) Confocal images showing viral expressions and VGLUT1 immunostaining in dorsal CA3. DAPI (blue), GCaMP8s (green), mCherry (red), and VGLUT1 antibody (magenta). Scale bar: 1000 µm. (B) High-magnification views of ipsilateral (i) and contralateral (ii) CA3 from boxed regions in (A). Scale bar: 100 µm (C, D) Quantification of mCherry (Cre-dependent) and VGLUT1 fluorescence intensity in ipsilateral vs. contralateral CA3 showing localized Cre delivery (p = 0.02) and significantly reduced VGLUT1 expression (p = 0.001) paired Mann-Whitney U tests. (E) Comparison of GCaMP event amplitude (p = 0.60, n = 7 mice for pCA3 and control dCA3, n = 4 mice for VGLUT1-cKO dCA3) (F) Proportion of place cells in familiar and novel environments. Both control and VGLUT1-cKO dCA3 displayed a higher proportion of place cells compared to pCA3, across contexts (Kruskal– Wallis H tests, familiar: H = 8.664, p = 0.013; novel: H = 10.929, p = 0.004; Dunn’s post hoc tests, familiar: p = 0.027 and 0.025; novel: p = 0.008; n = 7 mice for pCA3 and control dCA3; n = 4 mice for VGLUT1-cKO dCA3).

**Supplementary Figure 8.**
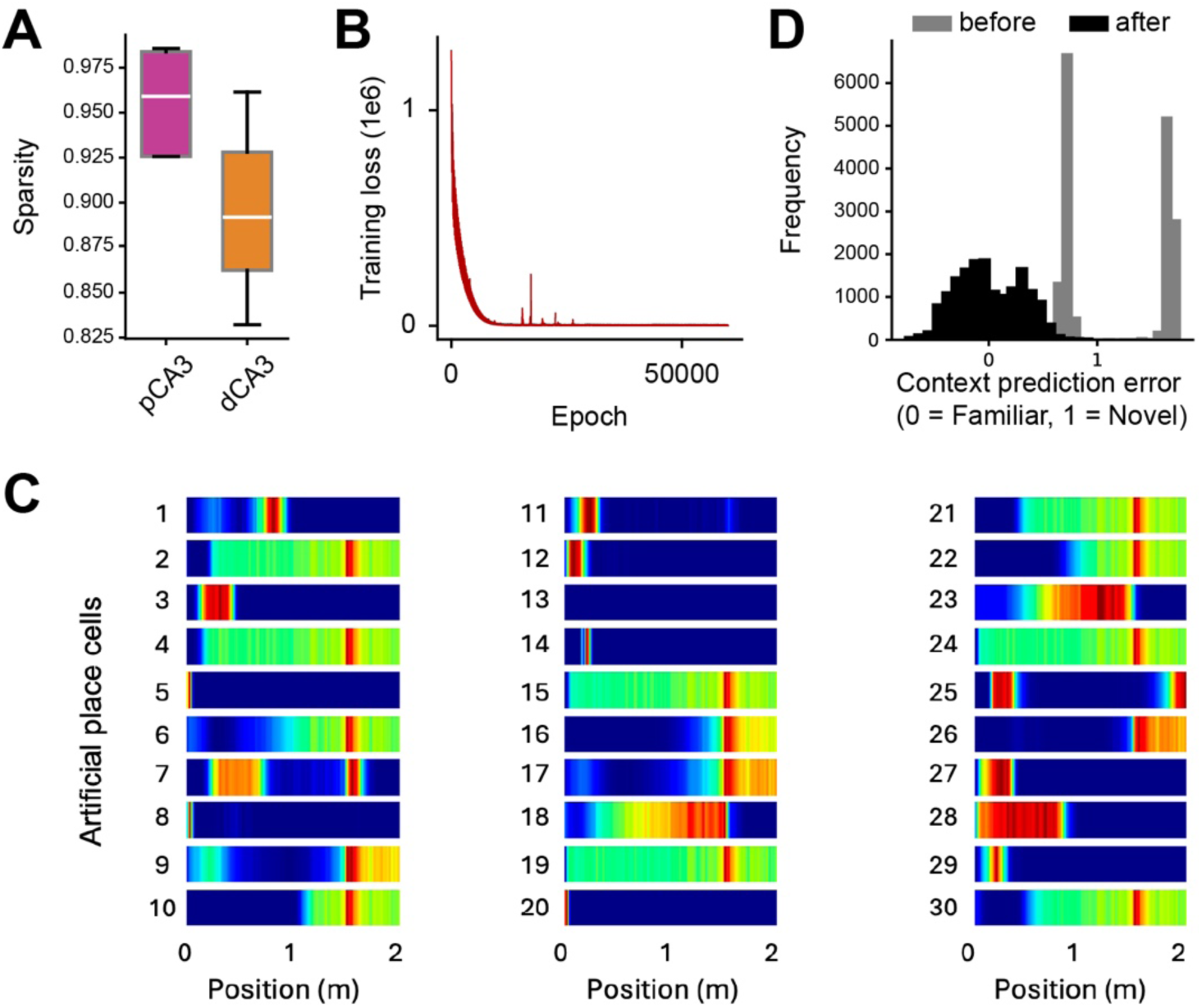
(Related to Figure 4) Spatial tuning properties of *in silico* place cells in RNN. (A) Comparing sparsity level via functional connectivity matrices: sparsity is evaluated based on the functional connectivity matrix of neurons in each subregion, using a 20-neuron window for both distal and proximal recordings. Results averaged across a recording session align with physiological studies, indicating greater recurrence in the dCA3 subregion. (B) Training loss of multi-task RNN: the training loss curve for the multi-task RNN (referenced in Fig. 4F) demonstrates the stability of the model’s convergence. (C) spatial RNN tuning curve with different neuron number: Example of tuning curves for a spatial RNN network, modeled with varying neuron counts (30 neurons), highlighting the network’s spatial encoding capabilities. (D) Distribution of prediction error across all laps before (gray) and after (black) training RNN.

**Supplementary Figure 9.**
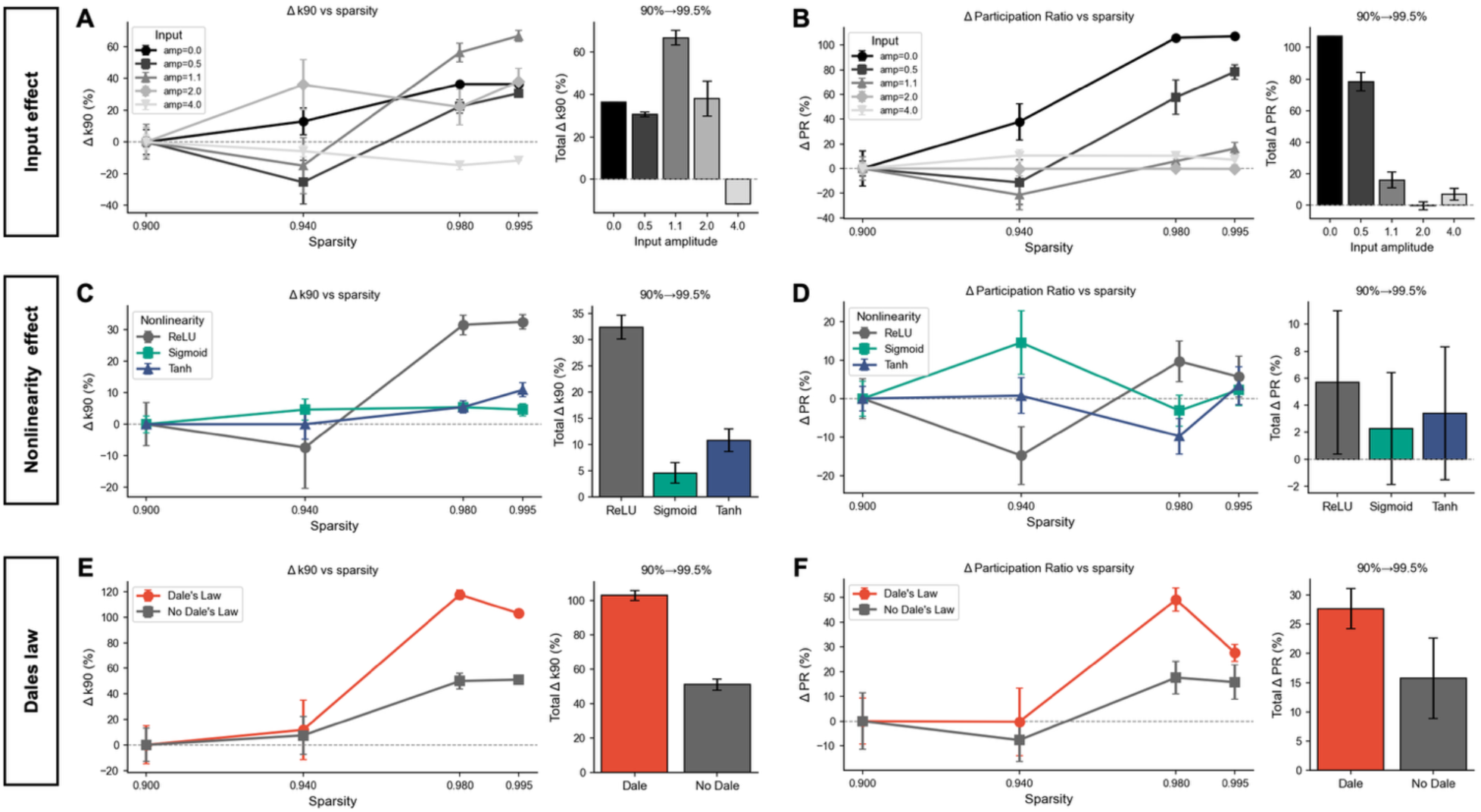
(Related to Figure 4, 5) Robustness of sparsity-driven increases in neural manifold dimensionality across biologically plausible recurrent network conditions. (A, B) Effect of input amplitude on the relationship between network sparsity and neural manifold dimensionality. Increasing recurrent connectivity sparsity from 90% to 99.5% results in increased dimensionality measured by k90 (A; number of principal components explaining 90% of variance) and Participation Ratio (PR; B), across input regimes ranging from noise-driven (amp = 0) to strongly input-driven (amp = 4.0). Line plots show changes in dimensionality as a function of sparsity; bar plots summarize the total change from 90% to 99.5% sparsity. (C, D) Effect of activation function. The sparsity–dimensionality relationship is preserved across different nonlinearities (ReLU, Sigmoid, and Tanh) for both k90 (C) and Participation Ratio (D). (E, F) Effect of Dale’s law. Networks constrained to obey Dale’s law (80% excitatory, 20% inhibitory neurons) exhibit sparsity-dependent increases in k90 (E) and Participation Ratio (F) comparable to networks without connectivity sign constraints. Across all conditions, increased sparsity consistently leads to higher neural manifold dimensionality. Error bars indicate SEM across independent network realizations (n = 10). Statistical significance for total changes was assessed using one-sample t-tests (*p < 0.05, **p < 0.01, ***p < 0.001). All simulations were obtained from recurrent neural networks (RNNs; N = 200 units) with randomly initialized sparse connectivity, where nonzero weights were drawn from a Gaussian distribution (σ = 1/√N) and normalized to a fixed spectral radius (ρ = 0.95).

**Supplementary Figure 10.**
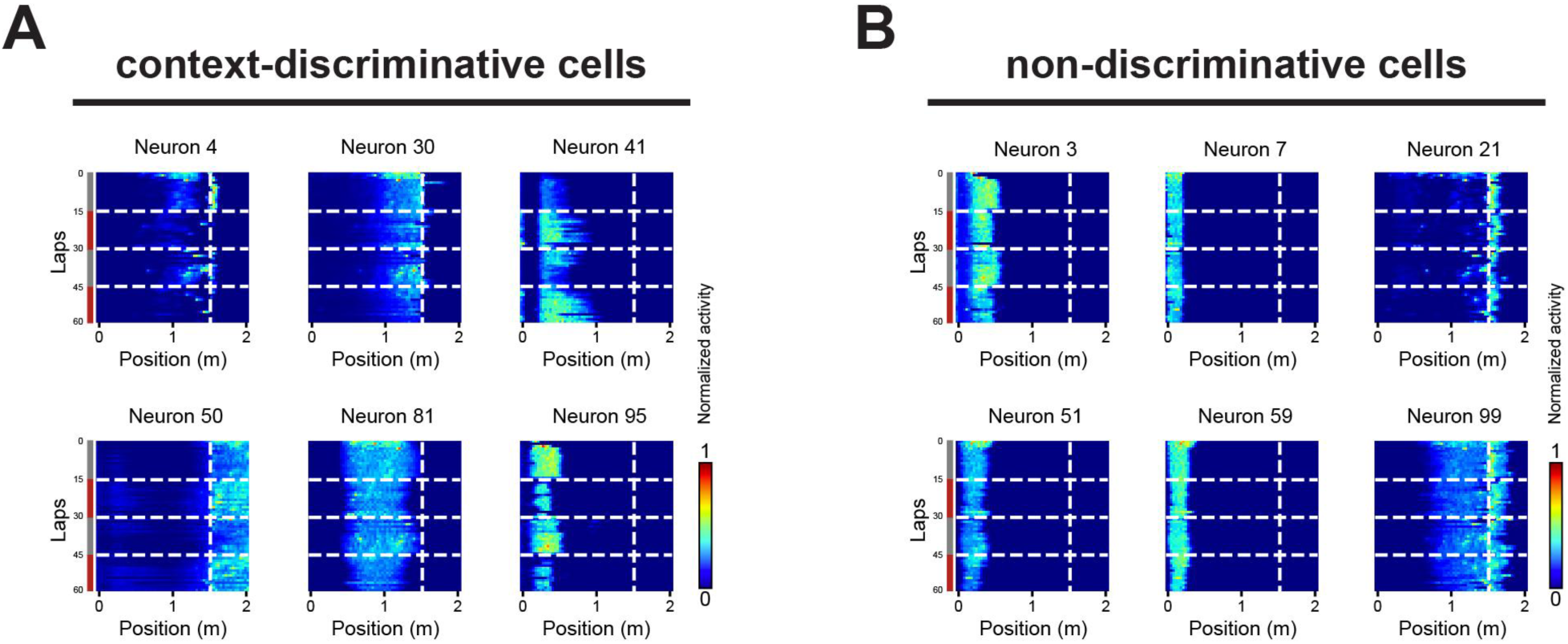
(Related to Figure 4) RNN artificial cell responses during context-switching task. Example heatmaps of lap-by-lap activity for RNN artificial cells are shown. Context-discriminative cells (A) and non-discriminative cells (B). Horizonal dashed lines indicate context changes, and vertical dashed lines indicate the reward location.

**Supplementary Figure 11.**
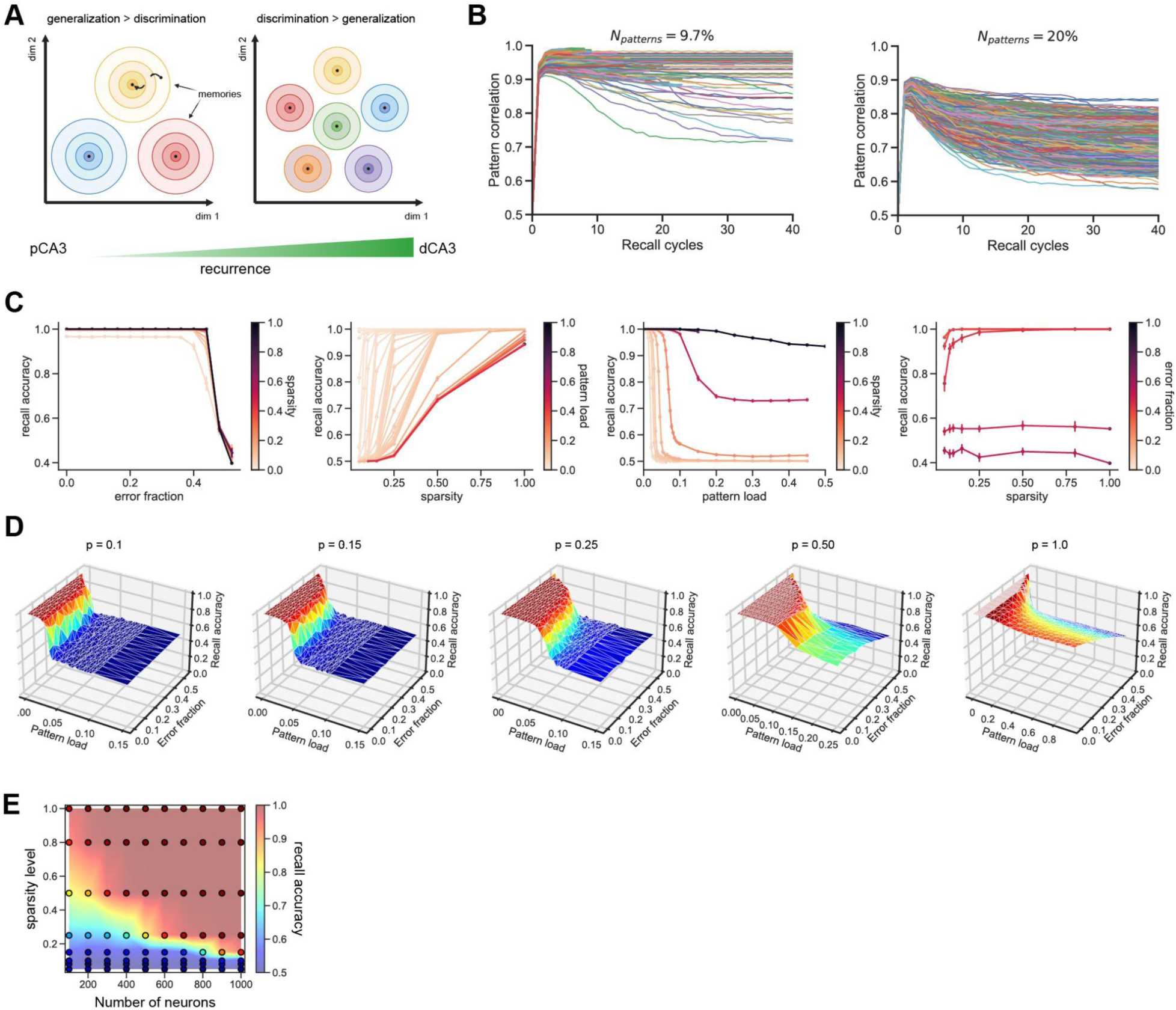
(Related to Figure 4) Hopfield model learning dynamics at different sparsity levels. (A) Schematic of energy landscape of sparse (pCA3) and dense (dCA3) network. The sparse network consists of fewer, larger energy minima to support generalization, while the dense network consists of more, smaller energy minima to support discrimination. (B) Learning dynamics at different pattern loads (*N*_*patterns*_: number of memories stored as a fraction of network size, here 1000 neurons; 1 line=1 simulation). Quality of recall is worse as more patterns are stored. (C) Pairwise relationships between parameters of interest (error fraction, sparsity, and pattern load) and recall accuracy, holding other parameters constant. (D) Dependence of recall accuracy on pattern load, error fraction and sparsity level. Sparser networks are able to store fewer patterns, but maintain robust generalization performance (high recall accuracy across range of input perturbations). Denser networks are able to discriminate more patterns, but do not generalize as well at higher pattern loads. In the discrimination (high pattern load, low error) regime, dense networks outperform sparse networks on a per-synapse basis. In the generalization (low pattern load, high error) regime, sparse networks outperform dense networks on a per-synapse basis (E) Neuron-synapse trade-off. Recall accuracy (color) for networks of different sizes (x-axis) and sparsities (y-axis) given a fixed number of patterns and error fraction = 0. Each point represents the result of one simulation; background: interpolated recall. Small dense networks have the same memory capacity as large, sparse networks, and both lie on the efficient frontier. This suggests a trade-off between increasing memory capacity by increasing neuron number and increasing synaptic density. Heuristically, the brain tends to be liberal with neurons but economical with synapses: memories are distributed across large, sparsely connected networks rather than small densely connected ones.

**Supplementary Figure 12.**
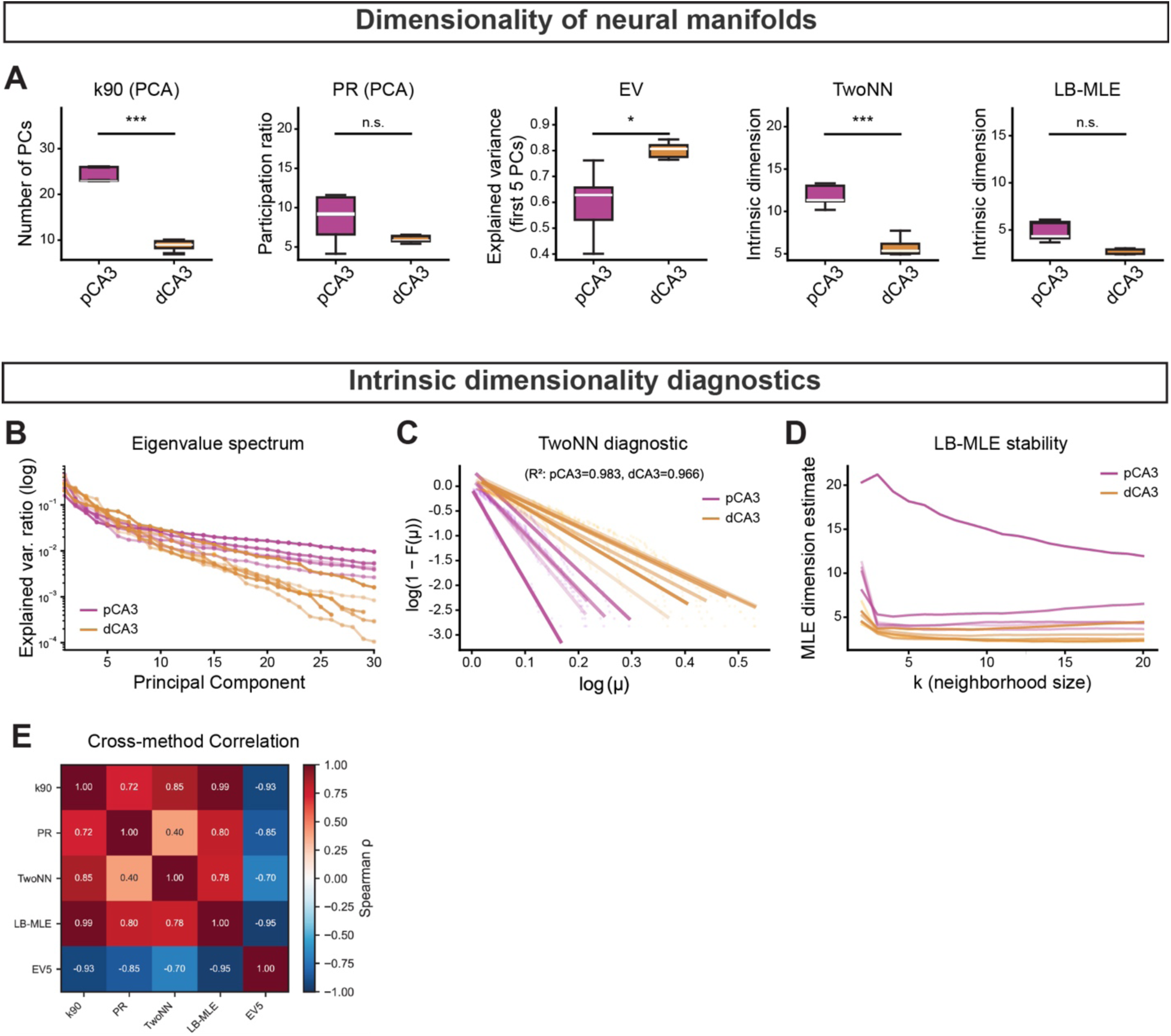
(Related to Figure 5) Dimensionality of neural manifolds in proximodistal CA3 and intrinsic dimensionality diagnostics. (A) From left to right: k90, participation ratio (PR), cumulative explained variance captured by the first 5 PCs (EV; inversely correlated with the dimensionality), TwoNN intrinsic dimension^100^, LB-MLE (Levina–Bickel) method for estimation of intrinsic dimensionality (see Methods). All the methods show the trend of higher dimensionality for pCA3 data. (B) Eigenvalue spectrum on a log scale; the decay rate reflects manifold complexity; pCA3 spectra decay more slowly, indicating higher-dimensional structure. (C) TwoNN diagnostic: log(1−F(μ)) vs log(μ), where μ = d₂/d₁ is the ratio of 2nd- to 1st-nearest-neighbor distances. Linearity validates the TwoNN model assumption; slope = −d. Mean R² values are reported per region. (C) LB-MLE dimension estimate as a function of neighborhood size k. Stable plateaus indicate robust estimates. (E) Spearman correlation heatmap across all five dimensionality measures (k90, PR, TwoNN, LB-MLE, EV5), confirming cross-method agreement for dimensionality differences trend. Note that EV5 is inversely related to the other measures (more variance in fewer PCs implies lower dimensionality). Mann–Whitney U tests were used to determine statistical significance unless otherwise stated. Colors are matched with purple for pCA3 and orange for dCA3. n.s. = non-significant, *p < 0.05, **p < 0.01, ***p < 0.001.

**Supplementary Figure 13.**
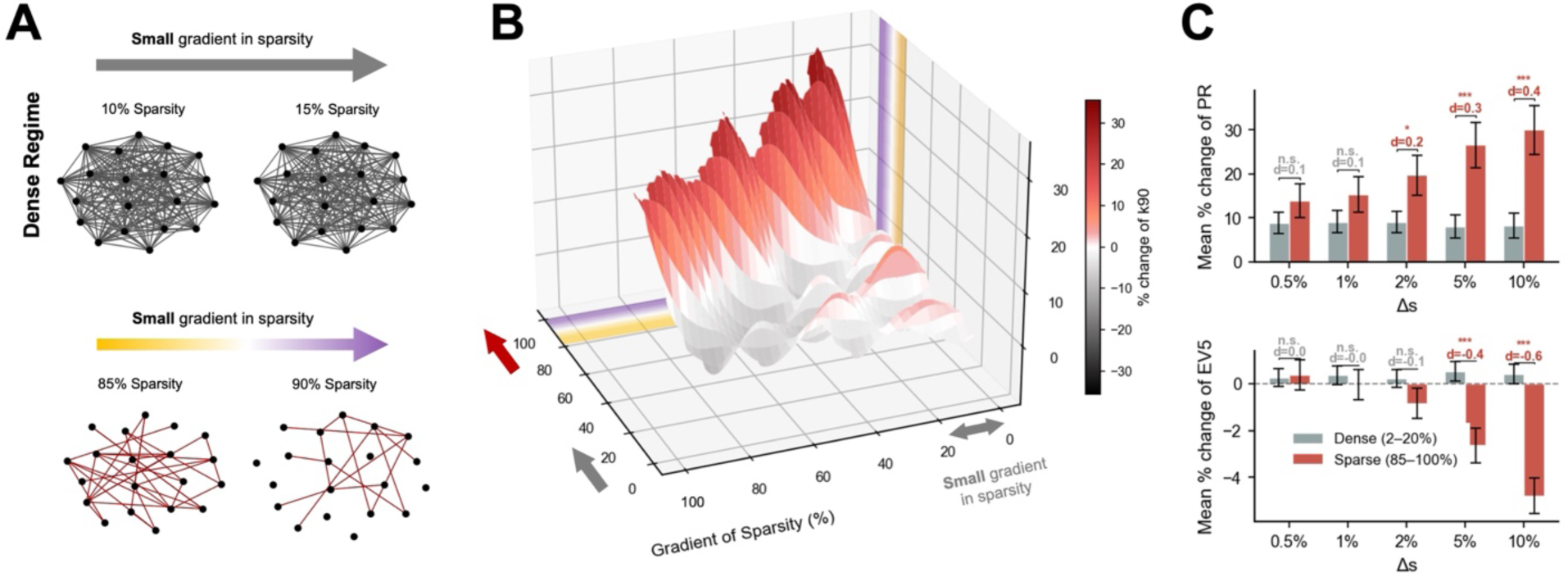
(Related to Figure 4, 5) Small changes in sparsity cause significantly larger shifts in the dimensionality of neural activity in sparse-regime networks compared to dense-regime networks. (A) Illustration of how a small, identical increase in sparsity reshapes connectivity differently depending on the operating regime. In the dense regime (top), raising sparsity from 10% to 15% removes only a handful of connections from an already richly connected graph, leaving the overall topology virtually unchanged. In the sparse regime (bottom), the same 5-percentage-point increase (from 85% to 90%) eliminates a substantial fraction of the few remaining connections, fundamentally altering the network’s communicative structure. Node positions are held fixed across all four graphs to facilitate visual comparison (*N* = 20 neurons). (B) Systematic exploration of how all possible combinations of average sparsity and sparsity gradient jointly determine the resulting change in manifold dimensionality. The surface displays the percentage change in k90 computed across every valid pair of sparsity levels in the fine-grained sweep. At low average sparsity (Dense regimes), even large gradients produce only modest changes in dimensionality, and the surface remains flat. As average sparsity increases, the surface rises steeply into a pronounced ridge, revealing that networks operating in the sparse regime amplify even small connectivity perturbations into large-scale reorganizations of the activity manifold. Color transitions from gray/black (dimensionality decrease) through white (no change) to red/dark-red (dimensionality increase). (C) Direct quantification of regime-dependent sensitivity. For each of five sparsity gradients (*Δs* = 0.5%, 1%, 2%, 5%, 10%), the signed percentage change in Participation Ratio (PR, top) and explained variance by 5 PCs (EV5, bottom) was computed separately for pairs whose midpoint falls in the dense regime (2–20%, gray bars) and the sparse regime (85–100%, red bars). At the smallest gradients (*Δs* ≤ 1%), dense and sparse networks respond similarly and no significant difference is detected. As *Δs* grows, sparse-regime networks diverge sharply: PR increases are several-fold larger than in the dense regime, while EV5 decreases more steeply; indicating that the activity spreads across more dimensions and becomes less compressible. This divergence is statistically confirmed by permutation test accompanied by growing Cohen’s *d* effect sizes (annotated in red), providing quantitative evidence that the sparse regime sits on a steeper slope of the sparsity–dimensionality landscape, where small connectivity changes have outsized functional consequences. n.s. = non-significant, *p < 0.05, **p < 0.01, ***p < 0.001.

## Key Resources Table

| REAGENT or RESOURCE | SOURCE | IDENTIFIER |
| --- | --- | --- |
| **Antibodies** |  |  |
| Rabbit anti-PCP4, 1:500 | Sigma-Aldrich | Cat. HPA005792;<br>RRID: AB_1855086 |
| Rabbit anti-VGLUT1<br>conjugated with Cy5, 1:500 | Synaptic Systems | Cat. #135303C5;<br>RRID: AB_2744598 |
| Anti-rabbit Alexa 647, 1:800 | Jackson<br>ImmunoResearch | Cat. #711-605-152;<br>RRID: AB_2492288 |
| **Experimental models: Organisms/strains** |  |  |
| Grik4-cre | Jackson Laboratory | Strain 006474 |
| *Slc17a7<sup>flox/flox</sup>* | Souter et al. 2021 | N/A |
| **Bacterial and virus strains** |  |  |
| rAAV9-syn-FLEX-jGCaMP8s-WPRE | This paper | Addgene 162377-AAV9 |
| rAAV9-syn-jGCaMP8s-WPRE | This paper | Addgene 162374-AAV9 |
| rAAV1-hSyn-Cre-WPRE | This paper | Addgene 105553-AAV1 |
| rAAV1.hSyn.DIO.mCherry | This paper | Addgene 50459-AAV1 |
| **Chemicals, Peptides, and Recombinant Proteins** |  |  |
| PBS | Thermo Fisher | 7732-18-5 |
| PFA | Electron Microscopy<br>Sciences | 15714-S |
| Triton X-100 | Sigma-Aldrich | T8787-100ML |
| Normal Donkey serum | Jackson<br>ImmunoResearch | 017-000-121 |
| Aqua-Mount | Fisher Scientific | 14-390-5 |
| **Software and algorithms** |  |  |
| Python3.7 | https://www.python.org | N/A |
| Unity | https://unity.com/ | N/A |
| SIMA | Kaifosh et al., 2014 | https://github.com/losonczylab/sima |
| ImageJ | Schneider et al.<sup>7</sup> | https://imagej.nih.gov/ij/ |

## Methods

All experiments were conducted in accordance with NIH guidelines and with the approval of the Columbia University Institutional Animal Care and Use Committee. No statistical methods were used to predetermine sample sizes. The experiments were not randomized, and the investigators were not blinded to allocation during experiments and outcome assessment.

### Animals

Imaging experiments were performed with healthy, 2- to 4-month-old, heterozygous adult male and female Grik4-cre mice (Jackson Laboratory, 006474) and homozygous VGLUT1 mice (*Slc17a7^flox/flox^*)^27^ on a C57BL/6J background. Mice were group-housed under normal lighting conditions in a 12-hour light/dark cycle. Ad libitum water was provided until the beginning of training for the spatial navigation task.

### Viruses

For CA3 pyramidal cell imaging in Grik4-cre mice, experiments were performed by injecting a Cre-dependent rAAV expressing GCaMP8s under the control of the synapsin promoter (rAAV9-Syn-FLEX-jGCaMP8s-WPRE-SV40; Addgene #162377; titer, 2.3 ×10^13^viral genomes per ml). For VGLUT1-cKO experiments, a combination of rAAVs was injected to disrupt VGLUT1 expression and monitor neural activity: rAAV1-hSyn-Cre; Addgene #105553, rAAV1-hSyn-DIO-mCherry; Addgene #50459, rAAV9-Syn-jGCaMP8s-WPRE-SV40; Addgene #162374).

### Surgery

All surgical procedures were performed on Grik4-cre mice (2-4 months old) under isoflurane anesthesia (4% induction, 1.5% maintenance in 95% oxygen). Mice were placed on a stereotaxic surgery instrument (Kopf Instruments), and their body temperature was maintained using a heating pad. Meloxicam and bupivacaine were administered subcutaneously to reduce discomfort. Following a skin incision, a craniotomy was performed over the right hippocampus using a drill. A sterile glass capillary loaded with rAAV was attached to a Nanoject syringe (Drummond Scientific) and slowly lowered into the right hippocampus. The proximal and distal CA3 were targeted with two x–y coordinates, each consisting of two different injection sites separated in z: AP –1.46, ML –1.75, DV –2.1, –1.9 and AP –1.46, ML –1.3, DV –2.3, –2.1 relative to bregma, with 60 nl of virus injected at each DV location. The pipette was held for 5-10 minutes after the last injection and then slowly retracted from the brain. The skin was sutured, and the mice were allowed to recover for 4 days before the window/headpost implant.

For hippocampal window and headpost implantation, as described previously^23,31,32^, a 3-mm craniotomy was performed over the right anterior hippocampus, centered between the two injection coordinates. We then slowly aspirated the cortex overlying the right dorsal hippocampus and implanted a 3-mm glass-bottomed stainless-steel cannula (1.5mm height) for optical access, while maintaining the CA1 region intact. Subsequently, a titanium headpost with layers of dental cement was attached to the skull. The mice received 1.0 ml of saline subcutaneously and recovered in their homecage on the heating pad. All mice were monitored for 3 days of post-operative care until behavior training began.

### Behavioral paradigm

After a 14-day recovery period from implant surgery, mice were water-restricted to 85-90% of their original body weight and habituated to handling and head fixation. They were then exposed to a 2-meter-long linear virtual reality (VR) corridor that remained consistent during training and recording^26,32^. At the end of the environment, an inter-trial interval of 2 seconds with a blank screen was included, and the mouse was simply teleported to the next lap.

For the next 10-14 days, mice were trained to run through the virtual environment and lick for a water reward. In the habituation phase, a drop of water was given in a non-operant manner when the subject mice entered the reward zone (RZ). Additional water droplets were provided as they continued licking within a 30 cm area. In the training phase, the reward was given in an operant manner where the correct lick within the RZ could trigger the first drop of reward and subsequent droplets with additional licking. Mice were trained to run at least 30 laps in the environment. After recording during the GOL task, the context-switching task was performed the next day.

For context-switch experiments, we used a completely distinct set of visual cues along the 200 cm track as a novel environment. Each recording session consisted of four blocks: 15 laps in the Familiar context, 15 laps in the Novel context, 15 laps in the Familiar context, and 15 laps in the Novel context. The reward location remained fixed at 150 cm for both contexts. For subject mice used in both pCA3 and dCA3 imaging experiments, we used different visual cues for the novel environments, while the familiar context remained the same for both sequences.

### Histology

After the completion of imaging experiments, mice were transcardially perfused with 40 ml of ice-cold PBS (Thermo Fisher), followed by 40 ml of ice-cold 4% paraformaldehyde (PFA; Electron Microscopy Sciences). The brain was then kept submerged in 4% PFA overnight at 4 °C. On the next day, coronal hippocampal sections (thickness: 100 μm) were obtained using a vibratome (VT1200-S, Leica, Germany). After brief washing with 0.3% Triton X-100 dissolved in phosphate-buffered saline (PBST), brain slices were incubated in a blocking solution (5% normal donkey serum, 0.3% PBST) for two hours at room temperature.

For PCP4 immunostaining, slices were incubated with the primary antibody (rabbit anti-PCP4 diluted 1:500; HPA005792; Sigma-Aldrich) overnight at 4 °C. After the sections were washed three times for 15 minutes with 0.3% PBST at room temperature, they were incubated with the secondary antibody (Alexa Fluor 647-conjugated donkey anti-rabbit IgG antibody diluted 1:800; 711-605-152; Jackson ImmunoResearch) for 2 hours. For VGLUT1 immunostaining, slices were incubated with Cy5-conjugated primary antibody (rabbit anti-VGLUT1 diluted 1:500; 135 303C5; Synaptic Systems) overnight at 4 °C.

Slices were rinsed three times for 15 minutes with 0.3% PBST, stained with DAPI (1:1000 dilution in 0.3% PBST), and then mounted on microscope slides. Fluorescence imaging was performed using a Nikon Ti-E A1R laser-scanning confocal microscope with ×10/0.45-NA Plan Apo (Nikon) or ×20/0.75-NA Plan Apo (Nikon) objective lens. 3 μm Z-stack images were processed using NIS Elements software (Nikon) or Fiji (ImageJ).

### *In vivo* two-photon imaging

All imaging was conducted using a 2-photon 8 kHz resonant scanner (Bruker) and high-NA multiphoton objective (XLPLN25XWMP2, 25x/1-NA, Olympus). For excitation, we used a 920 nm femtosecond-pulsed laser (Chameleon Ultra II, Coherent). Pockels cells were used to regulate the power of the laser reaching the tissue. Green (GcaMP8s) fluorescence was collected through an emission cube filter set (HQ525/70 m-2p) to a GaAsP photomultiplier tube detector (Hamamatsu, 7422P-40). A custom dual-stage preamp (1.4 x 105 dB, Bruker) was used to amplify signals prior to digitization. All images were acquired with 512 x 512 pixel resolution at 30 Hz.

### Calcium imaging data preprocessing

The preprocessing steps for the raw fluorescence signal have been described elsewhere^56,64^. Briefly, the imaging data was motion corrected using the SIMA software package^101^. The time average of each imaged cell was manually inspected, and an ROI was hand-drawn over each cell using a data visualization server program developed in the lab.

To track the activity of the same CA3 PNs across days, we used cross-day ROI registration as described previously^64,102^. Briefly, ROI masks from the time-averaged frame of day n were transferred and aligned to those of day m, and each neuron was assigned a unique ID to allow longitudinal tracking across sessions (days 1, 3, 5, and 7). Fluorescence was extracted from each ROI using the FISSA software^103^ package to correct for neuropil contamination using six patches of 50% the size of the original ROI. For each resulting raw fluorescence trace, a baseline F was calculated by taking the first percentile in a rolling window of 30 s, and a ΔF/F trace was calculated. The ΔF/F trace for each cell was smoothed using an exponential filter, and all further analyses were performed on the resulting ΔF/F traces. We detected statistically significant transients as described previously^56^ to use for place field calculations. All further analyses were implemented using Python using custom-written scripts.

### Spatial tuning analysis

The virtual environment was divided into 100 evenly spaced bins (2 cm), which were then utilized to bin a histogram of each cell’s neuronal activity. Neuronal activity was filtered to include activity from when the animal was running above 3 cm/s and to exclude activity during the 2-sec teleportation at the end of the 2-m track. The spatial tuning curves were normalized for the animal’s occupancy and then smoothed with a Gaussian kernel (α = 6 cm) to obtain a smoothed activity estimate.

To detect place fields, we generated a baseline distribution of spatial tuning curves for each neuron by randomly and independently shifting the activity on each lap a random distance in a circular manner. Following this, we recalculated the smoothed, lap-averaged tuning curve as detailed above. We repeated this process a thousand times, determining the 95th percentile of baseline tuning values at each spatial bin, which served as the threshold for significant spatial tuning (p < 0.05). Areas of space where the true spatial tuning curve surpassed this baseline threshold were identified as potential PFs. In order to be classified as a place cell, ROIs needed to have at least 3 consecutive significant bins (6cm), but less than 25 consecutive bins (50cm). Moreover, in order to avoid spurious detection of significant bins, we set an additional criterion that there must be activity within the PF boundaries for a minimum of 20% of laps. For place cell sensitivity, representing the proportion of laps that had active events within the significant field, the number of laps that had at least 1 binarized event within the significantly detected fields was divided by the total number of laps. For place cell specificity, the total number of events within the significant field was divided by the total number of events observed within the lap. Then, it was averaged across all laps to have a single value for each ROI. If multiple fields were detected, they were computed separately and averaged across fields to have a single value for each ROI. For spatial information, it was calculated as described previously^104^.

### Tuning-curve correlation

To determine the similarity of spatial representations across days, we computed Pearson correlations between the tuning curves of each place cell for all pairs of sessions recorded on different days. Only cells that exhibited place fields on both days were included in the analysis.

### Population vector correlation

To assess the stability of spatial representations over days, we calculated the population vector (PV) correlation (Pearson correlation) between two sessions imaged on different days. Only cells defined as place cells on both days were included in the analysis. The diagonal values of the correlation matrix represent the correlation between corresponding spatial bins across different sessions. Specifically, each diagonal value indicates the correlation between the PV in a given spatial bin in one session and the PV in the same bin in another session. We averaged these diagonal correlation values across all spatial bins. This process was repeated for all pairs of sessions, and the correlation coefficient was quantified as a function of the day interval between sessions.

We also used PV correlations to quantify the similarity of the spatial code between and across contexts – F1 vs F2, N1 vs N2, and F vs N. Cells that were defined as place cells in at least one of the two blocks were included in this analysis. The same quantification process for correlation coefficients was applied

### Discrimination index

As described previously^105^, the discrimination index (DI) was calculated for each cell to estimate the response preference toward a given context. The DI was defined by the following formula:

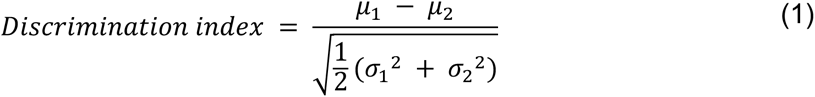

where μ and σ represent mean and standard deviations of calcium response amplitudes across laps of each environment, respectively^106^.

To test the significance of the DI value, we generated 1,000 surrogate data sets in which laps were randomly permuted and calculated the 95th percentile of the null value per cell. If the experimentally observed DI value surpassed the top or bottom 2.5% of the distribution of surrogate DI values, we considered it context-selective. Therefore, 5% of neurons were expected to have a significant DI value in a given session by chance.

### Population decoding analysis

To assess whether neuronal population activity patterns convey contextual information, we utilized a linear classifier support vector machine (SVM) to classify neuronal activity patterns into familiar or novel environments. For each imaging session, independent SVMs were trained and tested at each time point using a leave-one-trial-out cross-validation approach. Specifically, each decoder was trained with the neural population activity pattern from all laps except one withheld lap (n-1). We then evaluated whether the trained decoder accurately classified the held-out lap into the correct context. This process was repeated for each lap in the session, ensuring that every lap was utilized as a test lap at least once. Decoding accuracy was measured as the proportion of correct classifications. To compare the decoding efficacy between CA3 subregions, cells from proximal CA3 were subsampled to match the average detected ROI from distal CA3. The same SVM analysis procedures were then applied separately to the pCA3 and dCA3 datasets.

To further quantify context generalization independent of spatial position, we computed a cross-condition generalization performance (CCGP) as described previously^43,107^. A linear SVM decoder was trained on neural activity from one spatial bin (F–P1 vs N–P1) and tested on a non-overlapping spatial bin (F–P2 vs N–P2). This analysis evaluated whether population representations of context generalized across different spatial positions. CCGP values were averaged across all valid bin pairs within each session, and statistical comparisons between subregions were performed subsequently. In addition, we performed a position CCGP analysis to evaluate whether spatial representations generalized across contexts. In this analysis, the decoder was trained to discriminate two spatial locations (position A vs position B) in one context (familiar) and tested in the other (novel), and vice versa. Statistical comparisons between subregions were performed as previously.

To assess statistical significance relative to chance level, we generated a null distribution by randomly shuffling lap labels (context A or B; binary labels 0 or 1) 1,000 times and repeating the full decoding and cross-validation procedure for each shuffle. Empirical decoding accuracies from both the standard SVM and CCGP analyses were then compared to their corresponding shuffled distributions.

### Deep-superficial cell classification

To assess the functional heterogeneity of CA3 PNs along the superficial-deep axis, we performed principal component analysis (PCA) on the x and y coordinates of ROI centroids from distal CA3 datasets. The first principal component (PC1) was used to define a ‘middle line.’ The signed distance of each ROI centroid from this middle line was then calculated, with the sign indicating its relative position (positive for the top-right and negative for the bottom-left). These distances were normalized by the total spread of distances within each session To categorize ROIs into “superficial” and “deep” subclasses, the top 10% of ROIs with the highest normalized signed distances were classified as “superficial,” while the bottom 10% were classified as “deep.” The discrimination index (DI) was calculated for each group and compared to assess the functional differences between the superficial and deep PNs.

### Neural manifold analysis

We employed Principal Component Analysis (PCA) to reduce the dimensionality of neural data and visualize neural trajectories in a low-dimensional subspace. Neural activity was represented as a matrix *X* ∈ ℝ^*N*×*T*^, where rows correspond to ROIs (putative neurons) and columns to time samples. To ensure comparability between pCA3 and dCA3 regions, we randomly subsampled the number of neurons in pCA3 to match the average number of ROIs detected in dCA3 session. The data were mean-centered across time prior to PCA. We computed the neuron–neuron covariance matrix:

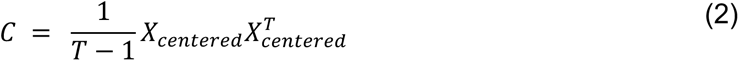

and performed eigen-decomposition of *C* to obtain eigenvalues {*λ*_ⅈ_}and corresponding eigenvectors (principal components). Data were projected onto the subspace spanned by the leading eigenvectors. Importantly, PCA was performed on time samples concatenated across trials/segments without prior trial-averaging, preserving trial-to-trial variability and the full population covariance structure and avoiding downward bias in dimensionality estimates.

To determine the dimensionality of the data, we identified the minimum number of principal components (k90) needed to explain at least 90% of the cumulative variance, calculated as:

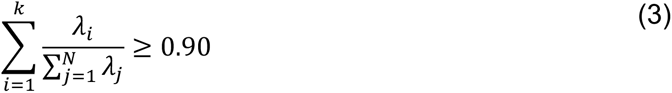

We also measured Participation Ratio (PR), which provides a continuous estimate of the effective dimensionality based on the distribution of all eigenvalues defined as

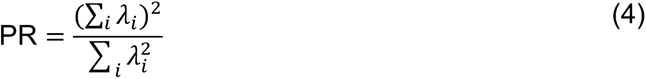

where *λ*_ⅈ_ are the eigenvalues of the covariance matrix (PR ranges from 1 for a single dominant component to *N* for a uniform spectrum); in addition to EV5, the cumulative variance explained by the first five principal components. The same manifold dimensionality quantification was applied to both neuronal recordings and sparse and dense recurrent neural networks (RNNs) from simulation.

### Intrinsic Dimensionality Estimation TwoNN intrinsic dimension

We estimated intrinsic dimensionality using the Two-Nearest-Neighbors (TwoNN) method^100^. For each data point, we computed the ratio *μ* = *d*_2_/*d*_1_ of the distance to the second nearest neighbor over the distance to the first. Under the assumption that data are locally uniformly distributed on a *d*-dimensional manifold, the empirical distribution of *μ* follows *P*(*μ* ≤ *t*) = 1 − *t*^-*d*^. The intrinsic dimension was estimated by linear regression of log (1 − *F*(*μ*)) on log (*μ*), where *F* is the empirical CDF, after trimming the extreme 5% tails of the distribution. The quality of the TwoNN fit was assessed via *R*^2^ of the log-log regression.

### Levina–Bickel MLE

We independently estimated intrinsic dimension using the maximum likelihood estimator of Levina & Bickel (2004)^108^. For a neighborhood of size *k*, the local dimension estimate at point *x*_ⅈ_ is:

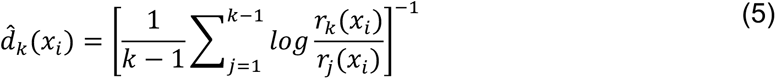

where *r*_j_(*x*_ⅈ_) is the distance from *x*_ⅈ_ to its *j*-th nearest neighbor. The global estimate was obtained by averaging 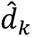 across all data points. To improve robustness, we averaged across four neighborhood sizes (*k* = 5, 10, 15, 20). Stability of the estimate was verified by plotting 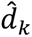 as a function of *k* for *k* = 2 to 20.

## Diagnostics and controls

To validate the estimators, we performed three controls: (1) eigenvalue spectra were plotted on a log scale to qualitatively assess the rate of spectral decay; (2) a cross-method Spearman correlation matrix was computed across all five measures (k90, PR, TwoNN, LB-MLE, EV5) to assess inter-method agreement. The dimensionality estimation of pCA3 and dCA3 subregions were compared using unpaired two-tailed *t*-tests and Mann– Whitney *U* tests for each dimensionality measure.

## Modeling

### Recurrent neural network modeling

Each subregion’s recurrent neural network (RNN) model consisted of a recurrent layer, alongside input and output layers. The network architecture and training method of our artificial neural networks was imitated from previous papers on path integrator artificial networks^109,110^. The recurrent layer comprised 100 rate neurons with tanh activation functions and was implemented using the TensorFlow LSTM package^111^. The output layer is a single (path integration) / two (multi-tasking network) neurons with dense connection from the recurrent layer with a linear activation function. The inputs to both the spatial imitator network and the multitasking network were the instantaneous speed and the initial position of the mice on the linear track. The initial position inputs were held constant, serving as the trial initiation signal. The network’s objective was to replicate the mice’s momentary position during each trial by integrating momentary speed and internally generated temporal signals within the network.

To stabilize network convergence, we introduced Gaussian noise as input to the network, a method shown to enhance convergence^112^. In the multitasking RNN (context switching + path integration), we included a noisy input signal simulating the context visual signal encoded as a binary code, combined with Gaussian noise centered around zero with a standard deviation of 0.3. This perceptual noise was added to mimic the context-dependent visual cue noise in the virtual arena. With a noiseless context signal, we anticipated a smaller neuronal population to be involved in context identification computations. The goal of the multitasking RNN was to minimize the Euclidean distance in a two-dimensional subspace between predicted location-context pairs and actual location-context pairs.

To implement recurrent sparsity, an *L1* regularization term was incorporated into the loss function, with different coefficients applied to promote either sparse (*λ*_*sparse*_= 0.1) or dense *λ*_*dense*_ = 0.001) connectivity. The total loss *L* is defined as:

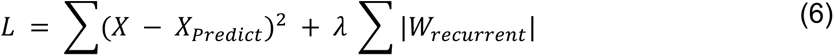

Where *X* denotes the vector of momentary location and context, *X*_*Pred*ⅈ*ct*_ is the network’s output, *W*_*recurrent*_ represents the recurrent weights across hidden layer neurons, and *λ* is the regularization coefficient.

Training was conducted using the Adam optimizer from the Keras package^111^, with a learning rate of 1e-4 and a minimum batch size of 8. We ensured the spatiality of training by verifying the saturation of the training loss function.

To reverse-engineer network activity, we analyzed the response functions of individual neurons in the hidden layer during a feedforward epoch, where instantaneous speed and context signals were provided as inputs. Analysis of the artificial neurons, including cell selectivity and population activity, followed the same criteria used for biological neurons in this study. Additionally, we replicated the formation of place-selective cells using a network with a smaller hidden layer size (30 neurons). Population-level results were based on simulations involving 100 hidden neurons.

### Hopfield networks

Hopfield networks were implemented in Python 3.8 in line with previous studies^113^, to model a CA3 subnetwork involved in storing and retrieving a set of memories at different levels of sparsity. In brief, a network was initialized with *N* binary valued neurons (for most simulations in this manuscript we use *N*=1000, but for a subset of simulations we vary *N* as an independent parameter as well, as indicated in the pertinent caption). For simplicity, we use the convention *ξ*_ⅈ_ ∈ {−1, +1}, which map onto the familiar ON (+1) and OFF (0), respectively. In the training phase, *M* patterns *ξ* are stored in the weight matrix according to the binary Hebbian rule

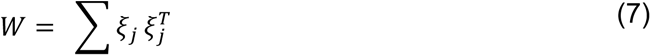

In the test phase, the trained network is initialized to a test pattern ξ’, constructed by flipping a predetermined fraction of the activities of a parent pattern ξ chosen from the *M* patterns the network was previously trained on. The network is then allowed to evolved according to the classical Hopfield dynamics

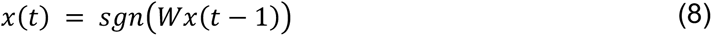

where *x*(0) = *ξ* until either convergence is reached (i.e., *x*(*t*) = *x*(*t* − 1)) or a maximum number of iterations *T*_max_ is reached.

To introduce sparsity into the Hopfield network, we choose a connection probability *p* and sample a structural connectivity matrix *A* where a connection exists between each pair of neurons i.i.d. with probability *p* (equivalently, a graph is drawn from the Erdős-Rényi process parameterized by *p*). The case *p=*1 reduces to the classical Hopfield network with full connectivity, while smaller *p* represents higher levels of sparsity. In sparse networks, weights between non-structurally connected neurons are not updated during training or used in the dynamics (which are otherwise unchanged), such that the effective weight matrix becomes

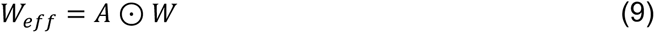

where ⊙ is the elementwise (Hadamard) product.

To model discrimination, we investigated how many distinct patterns networks at different levels of sparsity could store and accurately recall by drawing *M* random binary vectors for a range of *M* and storing them in the network. In the test phase, a fixed fraction of neurons were flipped and accuracy was read out as the similarity between the recovered pattern and the ground truth pattern as measured by Pearson correlation.

To model generalization, we investigated perturbations to a base pattern that could still be identified with the base pattern by networks at different levels of sparsity by fixing the number of patterns *M* and varying the fraction of flipped neurons. We again used similarity between the recovered pattern and the ground truth pattern as our accuracy metric.

### Sensitivity analysis of sparsity gradient across connectivity regimes

In order to measure and visualize the sensitivity of sparse regimes to small gradient of sparsity, we simulated a small rate recurrent neural network (RNN) with *N* = 10 units and ReLU activation. The network state evolved as:

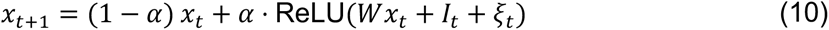

Recurrent weight matrices were generated by sampling entries *W*_ⅈj_ ∼ *N*(0, 1/√*N*) and independently zeroing each off-diagonal entry with probability equal to the target sparsity level *s* ∈ [0,1]. Diagonal entries were set to zero. Each matrix was then rescaled so that its spectral radius equaled 0.95. A two-dimensional sinusoidal input was provided to all units:

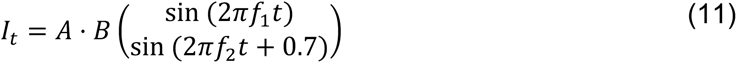

where *f*_1_ = 0.005, *f*_2_ = 0.011 cycles/step, amplitude *A* = 1.0, and *B* ∈ *R*^*N*×2^ with *B*_ⅈj_ ∼ *N*(0, 1/√2) was a fixed random projection matrix. Each network was simulated for *T* = 22,000 time steps. The first 200 steps were discarded as burn-in. Activity matrices *X* ∈ *R*^*N*×(*T*–200)^were mean-centered across time before analysis. Three PCA-based dimensionality metrics were computed from the mean-centered activity: (Participation Ratio (PR), k90, EV5).

#### Gradient surface construction

Sparsity was swept from 0% to 100% in 0.5% increments (201 levels). At each level, 10 independent networks were generated and simulated and per-repeat dimensionality measures were stored. For every ordered pair of sparsity levels (*s*_1_, *s*_2_) with *s*_2_ > *s*_1_ and *s*_1_ ≥ 2%, *s*_2_ ≤ 98%, the percentage change in each dimensionality measure was computed from repeat-averaged values as *Δm* = (*m*(*s*_2_) − *m*(*s*_1_))/*m*(*s*_1_) × 100. The resulting point cloud (indexed by average sparsity (*s*_1_ + *s*_2_)/2 and sparsity gradient *s*_2_ − *s*_1_) was interpolated onto a 120 × 120 grid using cubic interpolation and Gaussian-smoothed (*σ* = 3.0). Physically invalid regions (*s*_1_ < 2% or *s*_2_ > 98%) were masked (Supplementary Fig. 13B).

#### Dense vs. sparse regime comparison

To provide statical measure for the distinct effect of sparsity gradient in sparse and dense regimes, networks were classified by the midpoint of each sparsity pair into a dense regime (2–20% average sparsity) or a sparse regime (85–100%). For each gradient value (*Δs* ∈ {0.5%, 1%, 2%, 5%, 10%}), the signed percentage change was computed independently for each of the 10 repeats, preserving per-repeat variability. Group differences were assessed with two-sided permutation tests (5,000 shuffles), and effect sizes were quantified using Cohen’s *d* with pooled standard deviation (Supplementary Fig. 13C).

#### Connectivity visualization

For the schematic connectivity panel (Supplementary Fig.13A), weight matrices were generated with *N* = 20 neurons at sparsity levels of 10%, 15%, 85%, and 90%. Graph layouts used fixed node positions computed from a complete directed graph (spring layout, seed = 0), and edges with ∣ *W*_ⅈj_ ∣> 0.06 were displayed.

### Statistics

Statistical differences between means were determined by two-sided unpaired t-test and Mann-Whitney U test, Kruskal–Wallis tests with post hoc Dunn’s multiple comparison tests as mentioned in the text or figure legends. A p-value<0.05 was used as the criterion for statistical significance. Boxplots show the 25th, 50th (median), and 75th quartile ranges with the whiskers extending to 1.5 interquartile ranges below or above the 25th or 75th quartiles, respectively. All data analysis and visualization were done with custom software built on Python version 2.7.15, and 3.8.11. All data were expressed as mean ± SEM.

## Acknowledgments

We thank Bovey Y. Rao and Jingcheng Shi for comments on the early version of the manuscript, George Zakka for technical assistance, and Leo Rodriguez for the experimental schematic in Fig. 1B. Confocal imaging was performed with support from the Zuckerman Institute’s Cellular Imaging platform. T.G is supported by an R00MH129565 from the National Institute of Mental Health (NIMH). A.L. is supported by National Institute of Mental Health (NIMH) R01MH124047 and R01MH124867; National Institute on Aging (NIA) RF1AG080818; National Institute of Neurological Disorders and Stroke (NINDS) Brain Initiative U01NS115530; NINDS R01NS121106, NINDS R01NS131728, and NINDS Brain Initiative R01NS133381. E.K is supported by an overseas postdoctoral fellowship (RS-2024-00406980) from the National Research Foundation of Korea.

## Author Contributions

E.K., T.G., and A.L. conceived the project. E.K. and T.G designed experiments. E.K collected data, and performed analysis with assistance from T.G., T.S.M., and D.S.P.. E.Z. and Z.L. performed the computational modeling. All authors wrote the paper.

## Conflicts of Interest

The authors declare no competing interests.

## Notes

### Competing Interest Statement

The authors have declared no competing interest.

### Summary of Updates

new analysis supplementary figure added to the revision

